# First community challenge for automated virus taxonomy

**DOI:** 10.64898/2026.07.04.736517

**Authors:** Cédric Lood, Swapnil P. Doijad, Evelien M. Adriaenssens, Yiming Bao, Jakub Barylski, Ben Bolduc, George Bouras, J. Rodney Brister, C. Titus Brown, Antonio Pedro Camargo, Lander De Coninck, Sebastian Deorowicz, Robert Edgar, Robert A. Edwards, Shitao Gong, Arthur Gruber, Adam Gudys, Ernestina Hauptfeld, Anneliek ter Horst, Tianyang Huang, Jingzhe Jiang, Lars Kaderali, Jaebeom Kim, Mart Krupovic, Jens H. Kuhn, Elliot J. Lefkowitz, Mathieu Leobold, Shuai-Cheng Li, Yiyun Liu, F. A. Bastiaan von Meijenfeldt, Uri Neri, Judit J. Penzes, N. Tessa Pierce-Ward, Janina Rahlff, Alejandro Reyes Muñoz, Luisa Rubino, Sead Sabanadzovic, Jiayu Shang, Peter Simmonds, Martin Steinegger, Matthew B. Sullivan, Yanni Sun, Lili Tian, Yi-Gang Tong, Robert Turnbull, Dann Turner, Arvind Varsani, Ziye Wang, Yasas Wijesekara, Wytamma Wirth, Haolong Xia, Shuo Yang, TzeChing Yeo, Jinbei Zhang, Xianglilan Zhang, Shanfeng Zhu, Andrzej Zielezinski, Simon Roux, Bas E. Dutilh

## Abstract

The rapid rate of virus discovery renders manual curation by taxonomy experts increasingly impractical, creating a need for reliable software that can reproducibly assign viral contigs to taxa at all fifteen ranks of the virus taxonomy. We led an open community challenge for the computational taxonomic classification of viruses and assembled a dataset of virus sequences combining expert-curated and metagenomic sequences. Seventeen teams contributed a total of thirty-four automated, fully reproducible classification pipelines. Most tools correctly assigned viruses belonging to established species, genera, or families, but viruses that are unclassified at those lower ranks remain challenging. This study provides datasets, open-source software, novel approaches, and recommendations to benchmark computational taxonomic classification of viruses, and support organizing the many viruses discovered in big omics data.

## Introduction

Viruses dominate Earth’s biosphere in both abundance and genetic diversity, yet no single method can completely resolve their taxonomy. Taxonomy provides far more than names: it anchors inferences about host range, replication strategies, virulence potential, and ecological roles. This information is essential for interpreting viromics data, tracking emerging pathogens, designing phage therapies, and engineering viral vectors. There is consensus that viruses are so diverse that no single taxonomic method can be used to classify them all (Simmonds et al., 2023). Since its inception, the International Committee for the Taxonomy of Viruses (ICTV) has relied on the expertise of the global community of virologists to classify viruses into a ranked taxonomy (Gorbalenya, Krupovic, et al., 2020). This has generated a patchwork of methods and criteria that, ideally, capture the features of different viral lineages into meaningful taxa that reflect the biology and evolutionary relationships of viruses. This information is formalized in taxonomy proposals, written by expert groups, evaluated and ratified by the ICTV (Siddell et al., 2023). These documents provide evidence on how viruses within each taxon shall be classified, ideally based on specific demarcation criteria, and are available via the ICTV website at https://ictv.global/files/proposals/approved (Black et al., 2026).

Over the past decades, the exponential increase in DNA and RNA sequencing capacity and the related discovery of new viruses has brought new challenges. An important source of new viral genome sequences is metagenomics, i.e., the direct sequencing of genomic material from complex samples, also referred to as viromics when viruses are targeted specifically (Mokili et al., 2012; Simmonds et al., 2017). Viromics involves sampling, sample preparation, and assembly, which may result in incomplete or inaccurate sequences (Cook et al., 2024; Roux et al., 2019). The resulting viral sequences can be broadly organized into tens of millions of viral operational taxonomic units. Given the diverse methods that are currently used to classify viruses, it is unclear which approaches can be applied to metagenome-derived sequences. While viromics is rapidly expanding our view of the virosphere, researchers turn to taxonomy for structuring the resulting data (Dutilh et al., 2021), and for reasons of practicality and reproducibility, their taxonomic classification would ideally be automated.

There is currently no single ICTV-endorsed method for assigning viruses, often known only by their sequence, to established taxonomic ranks. Members of the ICTV Study Groups routinely use computational aids for taxonomy proposal writing, but a bioinformatics approach that enables researchers to compare the viral sequences representing viruses to those of all ICTV-established virus taxa and classify these viruses at an appropriate taxonomic rank is not available (Dutilh et al., 2021; Simmonds et al., 2023). An example for the challenges facing taxonomy is *Caudoviricetes*, a large class for bacterial and archaeal viruses. Currently, newly determined sequences that are ≥95% identical to sequences of already classified viruses are considered to represent viruses of the same caudoviricete species. Sequences that are ≥70% identical are considered to represent novel viruses requiring the establishment of novel species while being assignable to established genera (Turner et al., 2021). Implemented to accommodate the vast diversity of caudoviricetes, such universal, numerical demarcation criteria are uncommon in other viral clades, making the classification of novel or highly divergent viral sequences far from trivial. In the realm *Riboviria*, coronavirids are classified using pairwise distances between nodes on a phylogenetic tree derived from multiple sequence alignments of five concatenated protein domains (Gorbalenya et al., 2020; Woo et al., 2023). Further illustrations of the intricacy of classification criteria for viruses in the orders *Chitovirales* and *Picornavirales* are shown in Box 1AB.

The ICTV Computational Virus Taxonomy Challenge was designed to assess the urgent need for automatic taxonomic classification of new virus sequences by assessing different publicly available computational virus taxonomy tools or pipelines. We compiled a dataset of nearly 60,000 viral sequences and provided it online. Some of the sequences were generated specifically for the Challenge, derived from newly isolated and sequenced viruses with suggested classification by experts from ICTV Study Groups. Others were derived from virus genomic and metagenomic sequence repositories and edited to resemble genomic fragments that one might find in a metagenomic or viromic dataset. An open invitation was offered and broadly relayed online to any bioinformatics teams interested in participating in assigning these viral sequences using methods of their choice. An important requirement was that their pipelines were made available as open-source repositories, so that they might be readily used by others. Here we report the results, challenges, opportunities, and perspectives gathered during this first ICTV Taxonomy Challenge.

## Results

### Creation of a virus contig dataset for the taxonomy challenge

The dataset used for the ICTV Computational Virus Taxonomy Challenge was assembled from three separate sources. First, To verify whether the classification pipelines could correctly identify officially recognized taxa, we generated a dataset of 15,900 virus contigs representing 12,953 species from the ICTV Virus Metadata Resource (VMR38v3; Figure 1A). This total includes 617 contigs from non-viral mobile genetic elements that are within the remit of the ICTV (viroids, viriforms, and satellites). Second, to assess the performance of the pipelines in assigning novel viruses, we collected 840 previously uncharacterized virus contigs and invited ICTV Study Group members to suggest their classification (Figure 1B, see Methods). Except for two sequences associated with established caudoviricete genera (0.2%), these novel sequences were not assigned to established genera or species, but experts could assign them at the family or higher rank. The distribution across realms and kingdoms of these two datasets is shown in Figure 1C. Third, to challenge the various pipelines with unknown viruses, we included 3,151 predicted viral contigs from assembled metagenomic sequences available in the MGnify database (Richardson et al., 2023). The sequences were edited, as *de novo* sequencing and assembly commonly results in contigs with sequencing errors, or in fragmented or circularly permuted contigs (Figure 1ABD). This resulted in a dataset of 59,673 contigs, tailored to assess different strengths and weaknesses of the contributed pipelines. Each entry was assigned a random label, and the dataset was made available via GitHub (Supplementary Data 1). Participants were blinded to the sequence origins, classifications, and editing status for the duration of the challenge. This complete annotation for the dataset of 59,673 contigs can be found in Supplementary Table 1.

**Figure 1.**
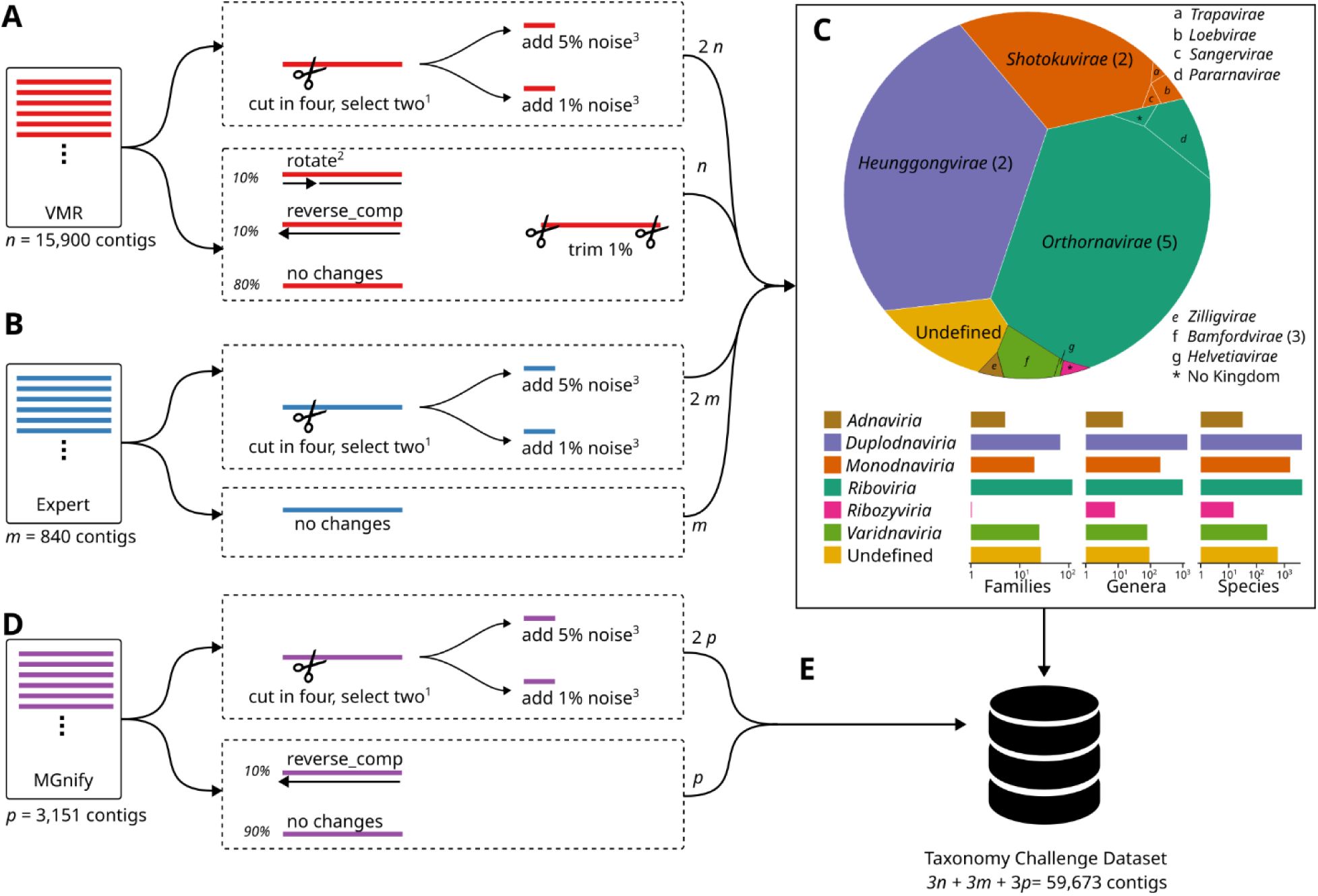
The dataset for the first ICTV Taxonomy Challenge consisted of 59,673 sequences that were derived from three sources as follows. **(A)** The ICTV Virus Metadata Resource (VMR38v3) was downloaded and all 15,900 sequences were edited by (i) cutting them into four equal parts, selecting two of the fragments at random then adding 1% or 5% random nucleotide substitutions, (ii) from 1—1,000 nucleotides of circular permutation and trimming 1% off the ends (10% of sequences), (iii) reverse complementing and trimming 1% off the ends (10% of sequences), or (iv) only trimming 1% off the ends (80% of sequences). **(B)** 840 diverse viral sequences were newly sequenced or identified in metagenomic datasets and assigned by experts from the respective ICTV Study Groups. These sequences were added either unchanged or cut (selecting two out of four fragments and adding 1% or 5% random nucleotide substitutions). **(C)** The sequences in A and B covered all realms defined in the virus taxonomy, specifically the Master Species List 39, version 4 (available at https://ictv.global/msl). Areas in the Voronoi plot represent the number of sequences contained in each realm. The labels correspond to the kingdoms within each realm, and the number of phyla within each kingdom is indicated between parentheses. **(D)** We identified viral contigs in assembled metagenomes from the MGnify database, selected 3,151 sequences, and added them unchanged (90% of sequences), reverse complemented (10% of sequences), or cut in four (selecting two out of four random fragments and adding 1% or 5% random nucleotide substitutions). **(E)** This composite dataset with 59,637 contigs was made available online for the challenge via the GitHub platform. <u>Legend</u>: ^1^selected at random, ^2^rotated at random from 1—1000 nt, ^3^nucleotide substitutions at random.

### Description of the entries

By the deadline for submission to the challenge (1^st^ January 2025), seventeen teams of bioinformaticians and virologists had entered a total of thirty-four separate sets of taxonomic classifications and made their pipelines publicly available via GitHub (Table 1). One parameter of the challenge was that participants use the Master Species List 39 version 4 (MSL39v4, available at https://ictv.global/msl) for assigning taxonomic labels, and an initial screen revealed that some of the entries had systematic encoding errors consisting of illegal strings for the taxa names at various ranks. Thus, we provided a validation script and gave all teams the opportunity to identify and fix any issues in their pipelines. We also encouraged the teams to improve the documentation of their tools in the form of dedicated “readme” files on the respective GitHub repositories (see also the Supplementary Methods containing extensive descriptions of each pipeline). These files were a required part of the challenge and used to assess reproducibility. The final classifications from all teams for all contigs can be found in Supplementary Data 2.

**Table 1:**
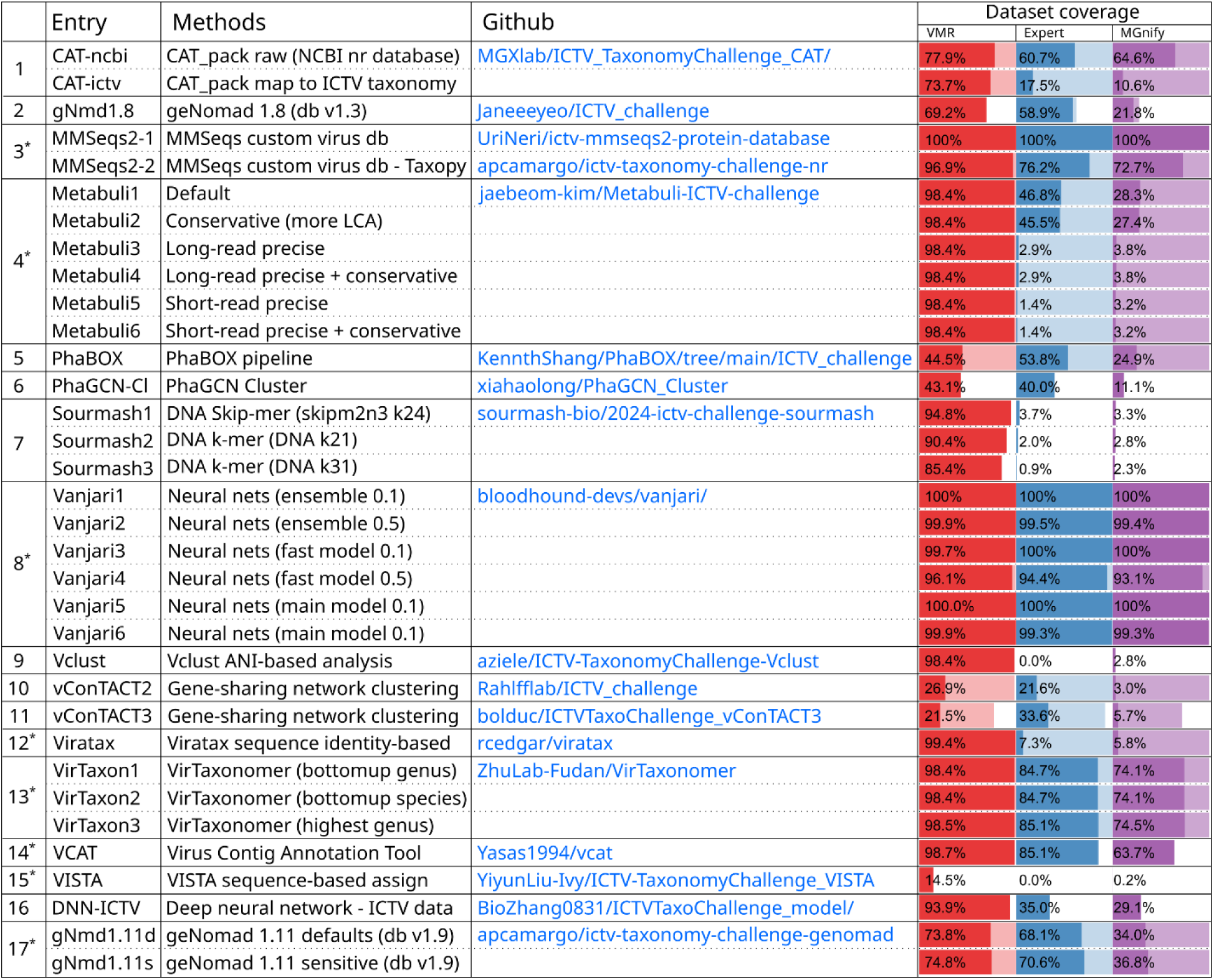
List of entries received for the ICTV Taxonomy Challenge. In total, 34 entries were received from 17 different teams. An asterisk in the first column indicates that confidence scores when available. The “Methods” column provides a high-level description of the pipeline (see Supplementary Methods for extensive description). All pipelines are publicly available on GitHub. The coverage columns indicate the fraction of sequences that were assigned a virus classification (solid color), Not Available (NA, transparent color), or completely missing (white color) in the VMR (red, n=47,700), Expert (blue, n=2,520), and MGnify (purple, n=9,453) datasets. Note that we distinguish pipelines without entries for a given sequence in the dataset (white color) and entries filled with NA (transparent).

Most final submissions resolved the issue of incorrect names within the ranks (Supplementary Table 2). The one clear outlier was CAT-ncbi, a pipeline consisting of contig annotation tool CAT (von Meijenfeldt et al., 2019) with its default NCBI database. Although NCBI implements ICTV taxonomy on all viruses in the database (Schoch et al., 2020), non-viral sequences are far more abundant, and include integrated proviruses that will be assigned to the taxonomy of their host. While CAT was proven to be efficient at classifying bacteria (Meyer et al., 2021), in the ICTV taxonomy challenge, CAT-ncbi assigned 31.5% of the viral sequences to non-viral taxa, highlighting the difficulty of annotating truly uncharted sequences, which might include both viral and non-viral contigs (Meyer et al., 2021). The remaining pipelines assigned valid taxa to the ranks used to assign the sequences, and most pipelines were error-free with five pipelines having further issues (incorrect fraction: 1.2%±2.6%). A separate analysis of the combinations of names across ranks (i.e., lineages) revealed that inconsistent lineage strings were common in ten pipelines, when evaluated based on the MSL39v4 (incorrect fraction:15.0%±21.6%, Supplementary Table 2).

Teams were not required to submit an assignment for all contigs but could also provide partial results. Eight teams submitted a total of 21 complete entries, whereas the remaining thirteen entries were partial, assigning between 14.5% (VISTA) and 99.9% (Sourmash) of the 59,673 sequences in the challenge dataset. VISTA (Zhang et al., 2025) was trained on complete genome sequences from 38 viral families and *Caudoviricetes*, and does not attempt taxonomic assignment for other sequences (see Supplementary Methods), limiting its scope and output. Sourmash (Irber et al., 2024; Pierce et al., 2019) uses sketching, storing only a subset of the *k*-mers in a sequence, which captures the overall sequence composition while reducing data size. This may miss short sequences that do not contain enough of the sketch *k*-mers or fall below the length threshold (≥200 nt). Among the pipelines that submitted incomplete entries, the fast nucleotide-based search tools Sourmash and Vclust (Zielezinski et al., 2025) delivered high sensitivity (94.8% and 98.4%, respectively) and low false-positive rates for sequences derived from known VMR viruses, but classified only a marginal number of entries from the Expert or MGnify datasets (Table 1). These tools are thus particularly suitable for rapid detection of viruses with high sequence similarity to those in the reference database. Some tools also included classification scores (marked with asterisks in Table 1), and changing such cutoffs may be valuable for sequences where classification is challenging, especially at higher taxonomic ranks.

### Accuracy for sequences associated with classified viruses

The VMR section of the challenge dataset included sequences from taxa of all major ranks of the ICTV taxonomy (Figure 1A) and was used to evaluate the classification accuracy for sequences derived from classified viruses or their close relatives. These query sequences should all be classifiable, given their high sequence similarity to reference viruses in the VMR. Figures 2A and 2B show the accuracy for all the VMR-derived sequences, and for the entries that were predicted by each pipeline, respectively. The latter excludes sequences that were not predicted, despite their close sequence similarity to reference viruses, and are represented by the white and transparent bars in the “Dataset coverage” section of Table 1.

**Figure 2.**
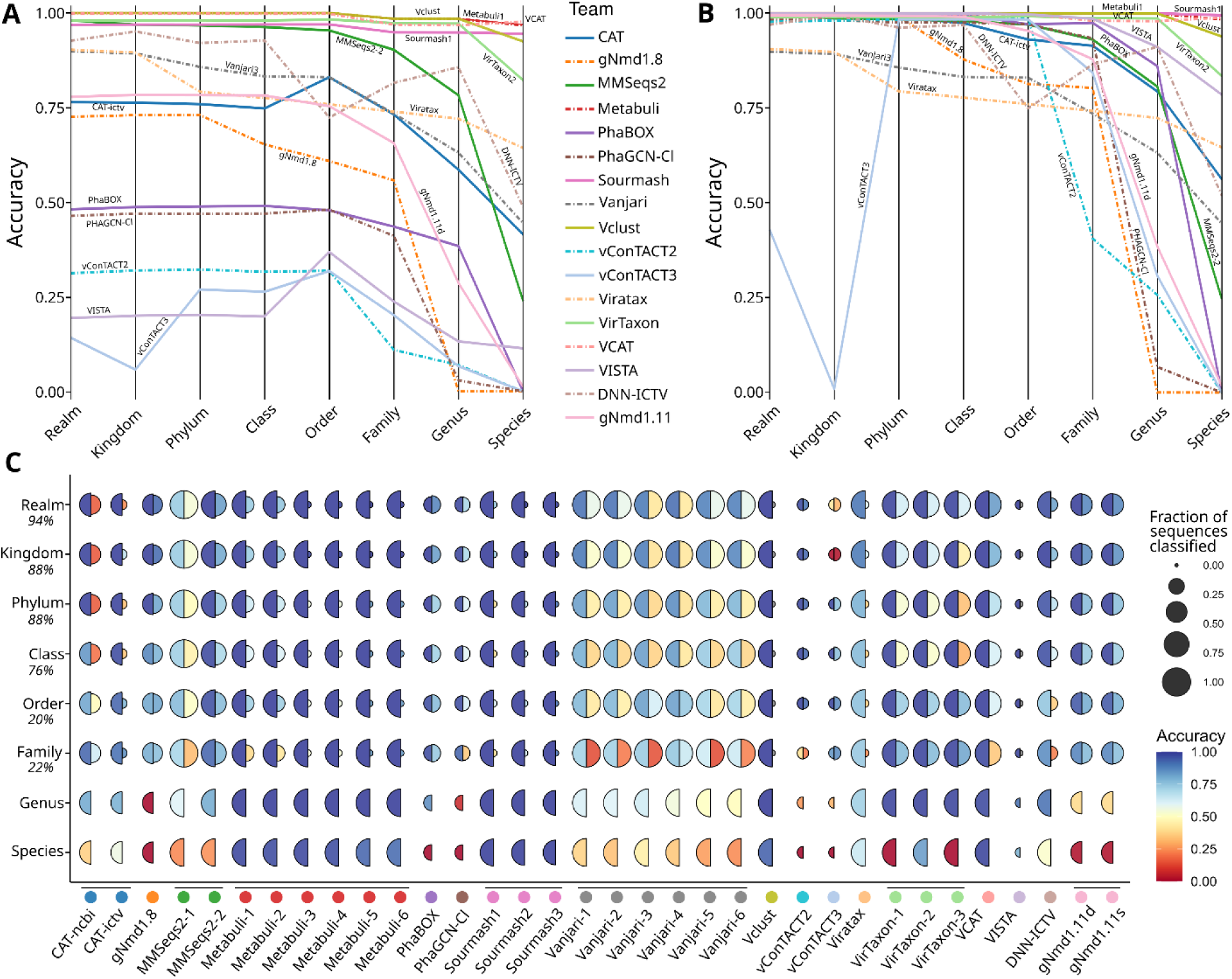
(A) Classification accuracy of the entries across the primary virus taxonomy ranks for sequences in the VMR datasets. Best-scoring contribution are shown when a team provided multiple entries(based on score for the species rank). (B) Classification accuracy restricted to the predicted entries, i.e., excluding empty, “NA”, or “unclassified” calls. (C) Split-bubble plot showing the fraction of sequences classified (radius) and classification accuracy (color) at each taxonomic rank for the VMR and Expert datasets (left and right half circles, respectively). As only 2/840 sequences were assigned below the family rank in the Expert dataset, the bottom two lines lack the right-hand half circles.

In general, most pipelines were more accurate at higher taxonomic ranks than at lower ones, indicating that sequences are commonly misassigned to different species within the same genus or family. For example, for geNomad, PhaGCN Cluster (Xia et al., 2026), and VISTA (Zhang et al., 2025) the accuracy decreased sharply below the family rank, whereas vConTACT2 (Bin Jang et al., 2019) dropped below the order rank. Similarly, PhaBOX (Shang et al., 2023) retained accuracy down to the genus rank and then decreased sharply. Vclust, Sourmash, and Metabuli (Kim & Steinegger, 2024) maintained a high accuracy across ranks with a slight decrease below the genus rank, whereas VCAT had a different pattern, as its accuracy marginally increased for species.

Next, we evaluated the classification performance on the VMR and Expert datasets separately (left and right half circles in Figure 2C, respectively). Within the VMR dataset, Metabuli, Sourmash, Vclust, VirTaxon-2, and VCAT assigned most sequences with high accuracy at all ranks down to species, whereas CAT-ICTV, geNomad, PhaBOX, PhaGCN Cluster, and VISTA assigned a subset of the sequences down to the family rank with high accuracy. Together, these results highlight the ability of these tools to detect similar sequences with up to 5% of random mutations, even when only a fragment was available.

For the VMR dataset, accuracy at higher taxonomic ranks may depend on the ability to trace the annotations of parent clades. As virus taxonomy is traditionally built bottom-up, i.e., beginning with species and genera, certain viruses may belong to “floating” taxa, i.e., taxa that have not been included into higher-ranking taxa. For instance, the most current ICTV taxonomy (Hendrickson et al., 2026) includes one floating genus *(Dinodnavirus),* 29 floating families, and a floating class (*Naldaviricetes*). None of these floating taxa could yet be reliably linked to any established realm, but it is also unclear whether they represent novel realms. Floating taxa may be connected to the virus taxonomy as it is further resolved. For example, the family *Anelloviridae*, floating in the MSL39v4, was assigned a full lineage in realm *Monodnaviria* in the MSL40v1. Note that *Monodnaviria* has been recently reorganized into four separate realms (Krupovic & Koonin, 2026). However, here we refer to the former taxon to remain consistent with the taxonomy at the time of the Challenge. Floating taxa have also been established when, for instance, realm membership is clear but the lower taxa are not. For instance, the realm *Riboviria* includes a floating family, *Tonesaviridae,* which could not yet confidently be assigned to an order, class, phylum, or kingdom.

In case of absent labels in the taxonomic lineage of a reference virus, pipelines that did not assign the sequence to any taxon were considered correct. As expected, all the pipelines predicted less accurately for the Expert dataset, which included novel sequences that were not yet present in the database (Supplementary Table 3). For the upper ranks from Realm to Class, to which the Study Group experts assigned most contigs (76% -> 94%, Figure 2C), geNomad, MMSeqs2-2, PhaBOX, PhaGCN Cluster, vConTACT2, VCAT, Metabuli, VirTaxon1-2, Sourmash2-3, and DNN-ICTV maintained an accuracy >50%. For the order and family ranks, a lack of expert annotation (20% and 22%, respectively) made the data difficult to interpret.

### Consistent classifications across sequence transformations and pipeline configurations

One of the aims of the ICTV Computational Virus Taxonomy Challenge was to identify the strengths and weaknesses of different pipelines in the face of *de novo* assembled contig sequences such as may be obtained from metagenomics. To this end, sequences were edited in different ways, including the introduction of single nucleotide replacements to reflect sequencing errors or substitution mutations, fragmentation of viral genome sequences into shorter contigs, reverse complementation, and circular permutation of the viral sequences to mimic results from *de novo* assembly (Figure 1ABD). This was used to assess how the classification accuracies of all the pipelines are impacted by these transformations (Figure 3A, Supplementary Figure 1, Supplementary Table 4). A general emerging trend was a decrease in classification accuracy as transformations and random substitutions were introduced, but these effects varied among pipelines. Some pipelines, including Metabuli, Vclust, VCAT, geNomad, and VISTA, appeared to be resilient to such changes, whereas for PhaBOX and Vanjari, accuracy decreased with increasing substitution rate. Vanjari was strongly impacted by reverse complementation, an issue that has been addressed by the Vanjari team (Supplementary Methods), and PhaBOX2 was recently released that address some of these limitations (Shang et al., 2026).

**Figure 3:**
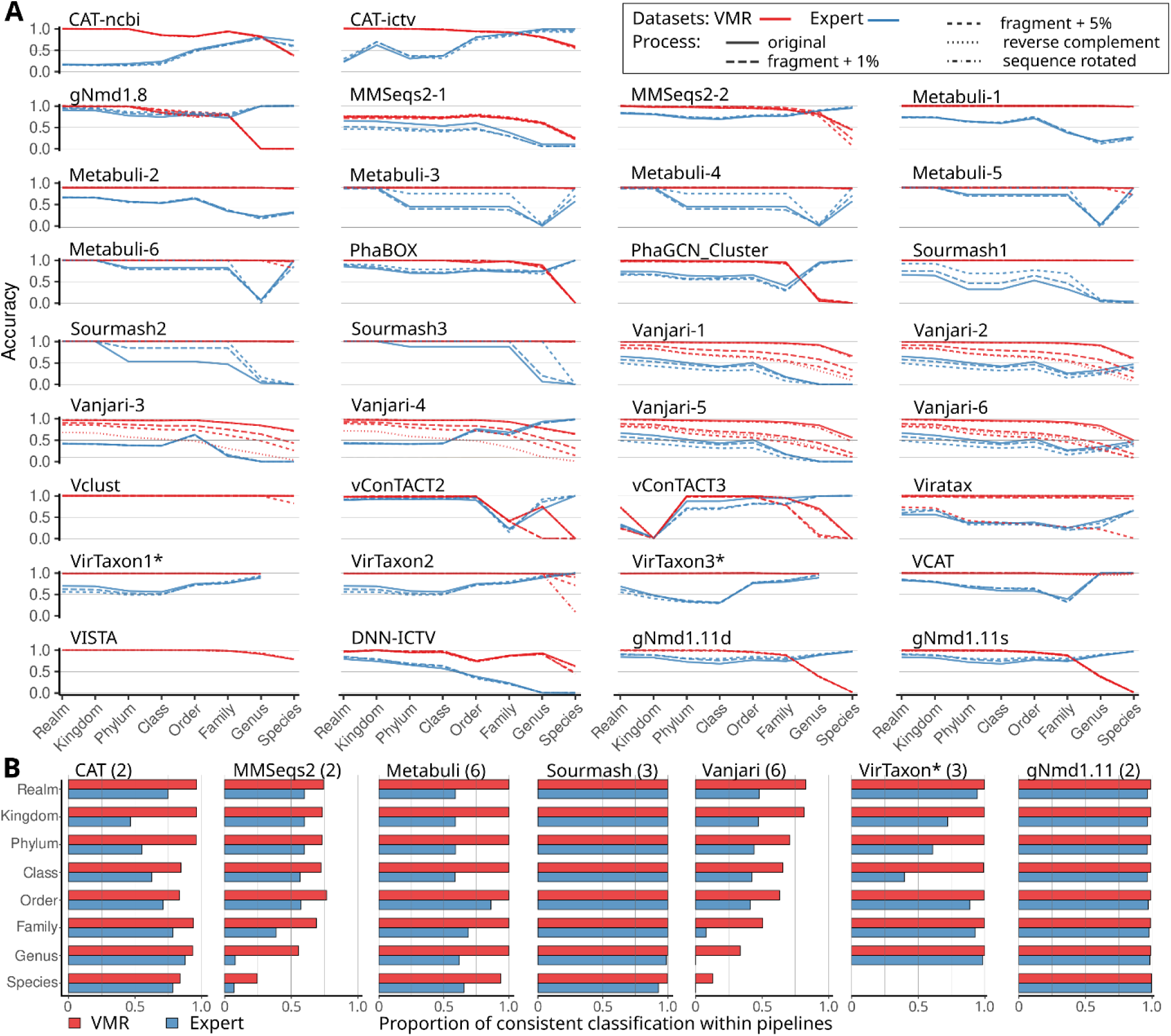
(**A**) The VMR and Expert datasets included sequences with known taxonomic classification that were edited by fragmentation, random nucleotide substitutions, reverse complementation, and circular permutation. Shown is how these various transformations impacted classification accuracy at all taxonomic ranks. Note that the VISTA and Vclust pipelines did not assign any sequences in the Expert dataset (Table 1). (*) VirTaxon1* and VirTaxon3* do not produce species rank predictions. (**B**) When multiple classifications were made by the same overall pipeline with different configurations (number indicated between parentheses after the pipeline name), the within-pipeline fraction of consistently predicted taxonomic labels was calculated to assess their consistency.

Several pipelines had high accuracy scores for genus and species classifications in the Expert dataset. As only two sequences were assigned below the family rank, these high scores reflect the fact that these pipelines did not assign most sequences at these ranks. While the lack of taxonomic assignment is considered correct in this particular case, such high accuracy values should still be interpreted with caution. Other pipelines, including MMSeqs2-1, Sourmash, Vanjari, and DNN-ICTV, did spuriously assign many sequences at the lowest two ranks that could not be ascertained by experts, and had lower accuracy scores as a direct result.

Seven pipelines were run with several different parameter settings (Metabuli, Vanjari, VirTaxon, geNomad 1.11, Sourmash) or different databases (CAT, MMSeqs2). In these cases, consistency among the multiple entries was assessed by calculating the proportion of sequences assigned consistent labels, including the empty label (or NA), across the variants of each of the pipelines, at the main ranks (Figure 3B). The most consistent pipelines were Sourmash and geNomad 1.11, assigning consistent labels to the sequences in both VMR and Expert datasets, while VirTaxon and Metabuli showed consistency within the VMR dataset. Finally, MMSeqs2 and Vanjari results had lower consistency within the lower ranking taxa.

### Classification performance across viral families

To assess how well different types of viruses could be computationally classified, we calculated classification metrics for each of the 278 virus families in our combined VMR and Expert datasets. The results are shown in Figure 4 for families in realms with RNA-based genomes, and in Figure 5 for DNA-based genomes and the families with an “Undefined” group (i.e., families not assigned to any realm), consisting of DNA-based viruses except *Pospiviroidae*, *Ovaliviridae*, *Halspiviridae*, and *Avsunviroidae*. The 18,258 sequences in “Undefined” families were not evaluated here, as they may not represent single taxonomic units. The number of available sequences varied widely among families, ranging from 47 families represented by three sequences each, to others with thousands (maximum: *Geminiviridae* with 2,814 sequences). The accuracy and false discovery rate (FDR) of each entry was separately assessed on sequences without and with mutations (paired heatmap columns in Figures 4 and 5). Some pipelines, such as PhaBOX, vConTACT2, and vConTACT3 differed considerably among realms, with good performance for *Duplodnaviria* (Figure 5) but poor performance for *Riboviria* and *Ribozyviria* (Figure 4). Although vConTACT2 was designed for the classification of prokaryotic viruses and this constraint was removed in vConTACT3, the pipeline, at the time of testing (though not in latest versions), did still require information about the domain of the host. As the default was set to a prokaryotic host, this complicated its testing under the conditions of the Taxonomy Challenge. Similarly, PhaBOX was mainly tailored to prokaryotic viruses, its successor PhaBOX2 has expanded its database and updated its underlying models to cover a broader viral diversity (Shang et al., 2026). Other pipelines, including Metabuli, Sourmash, Viratax, Virtaxon, VCAT, and DNN-ICTV, had similar accuracy scores for most families, with some exceptions discussed below. Depending on the pipeline, accuracy was associated with the number of reference sequences within a family, as reflected by the sequence count, with correlation values ranging from −0.2 for PhaBOX to 0.4 for Vanjari-4 (Supplementary Figure 3). The value was positive for most tools, showing that it was generally easier to classify viruses in large families than those in smaller ones.

**Figure 4:**
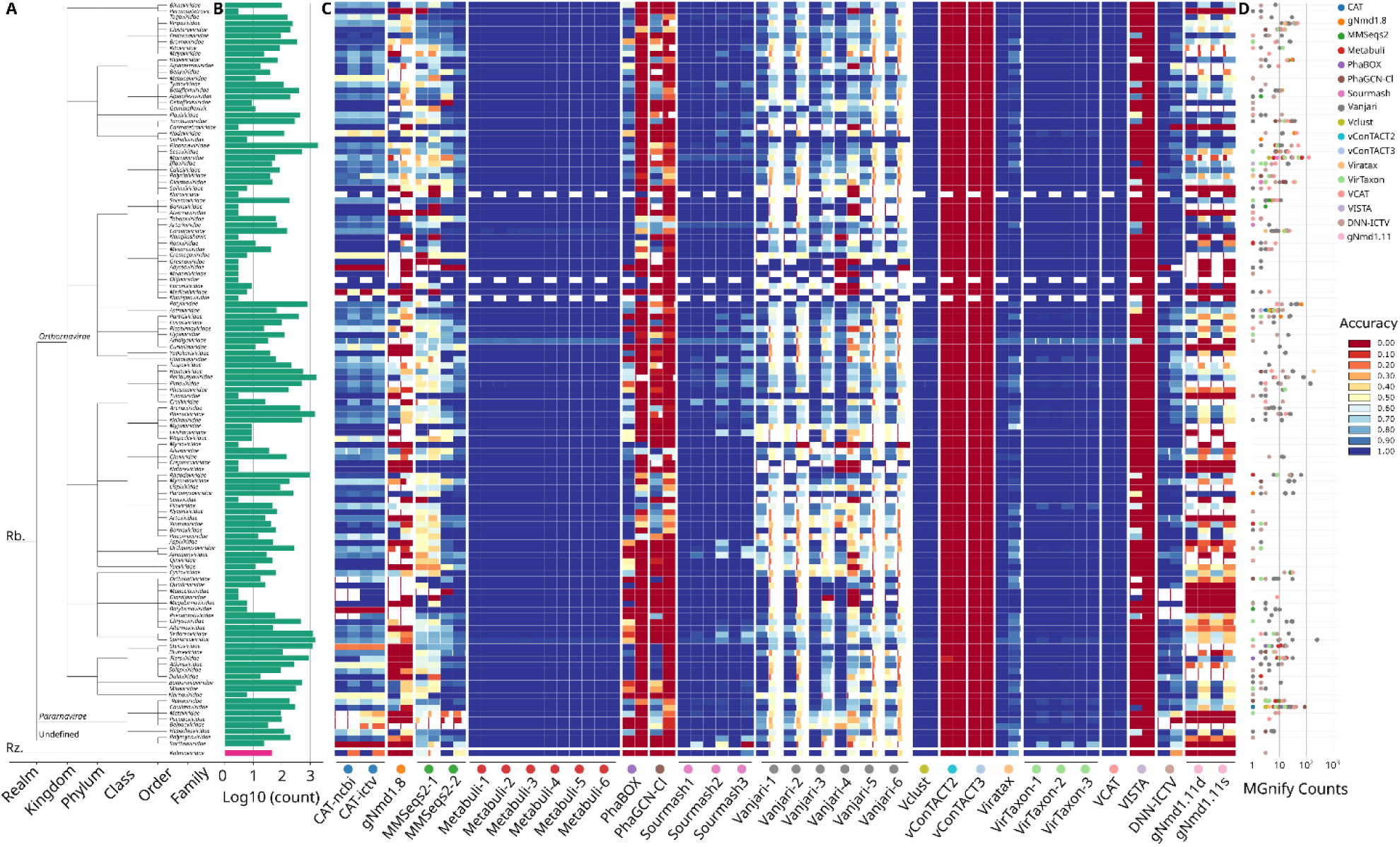
Results for all 122 taxonomic families of viruses with RNA genomes. **(A)** Cladogram showing families organized by realm, including Riboviria (Rb.) and the Ribozyviria (Rz.). **(B)** Bar chart showing the number of sequences in the VMR and Expert datasets combined (log scale). **(C)** Heatmap showing the accuracy (color) and inverse false discovery rate (tile width) on i) sequences without substitutions, rotations, or reverse complementation (left column for each entry) and ii) sequences that were cut and included 1% of random substitutions (right column). For 5% mutations, see Supplementary Figure 2. Floating families not included in higher ranks are not shown. **(D)** Number of sequences within the MGnify dataset that were assigned with high confidence (accuracy * (1 - FDR) > 0.9) into each family by each pipeline.

**Figure 5:**
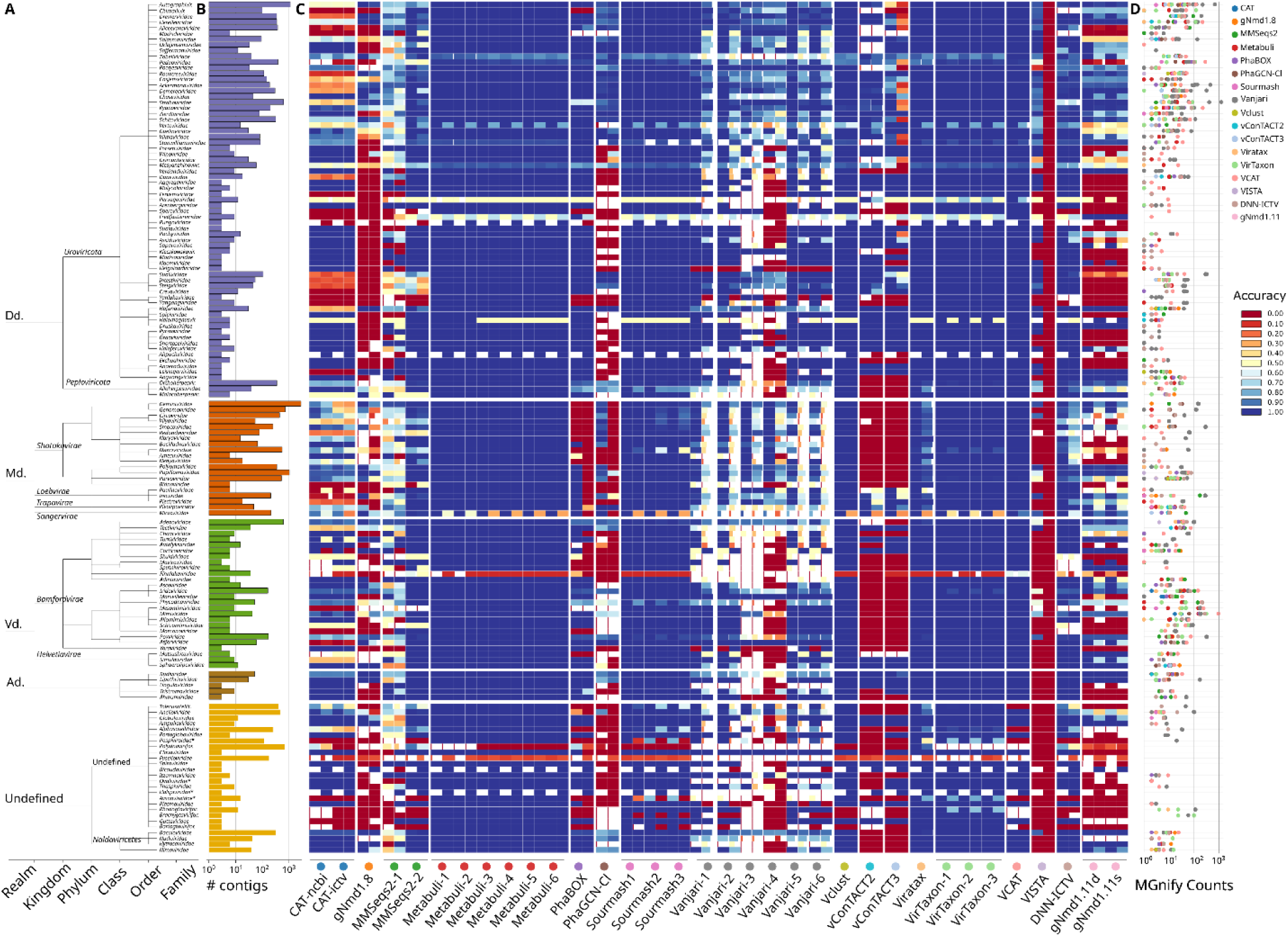
Results for the 152 taxonomic families for viruses with DNA genomes (+4 floating families with RNA genomes). **(A)** Cladogram showing families organized by realm, including Duplodnaviria (Dd.), Monodnaviria (Md.), Varidnaviria (Vd.), and Adnaviria (Ad.), and Undefined. Four RNA virus families are indicated with asterisks. **(B)** Bar chart showing the number of sequences in the VMR and Expert datasets combined (log scale). **(C)** Heatmap showing the accuracy (color) and inverse false discovery rate (tile width) on i) sequences without substitutions, rotations, or reverse complementation (left column for each entry) and ii) sequences that were cut and included 1% of random substitutions (right column). For 5% mutations, see Supplementary Figure 2. Floating families not included in higher ranks are not shown. **(D)** Number of sequences within the MGnify dataset that were assigned with high confidence (accuracy * (1 - FDR) > 0.9) into each family by each pipeline.

Analysis of the viruses with RNA-based genomes (Figure 4) revealed that for all families, sequences could be correctly assigned by at least one of the pipelines. Vclust, Sourmash, Viratax, VCAT, Metabuli, and VirTaxon assigned sequences to families correctly, even in the presence of random substitutions and fragmentation. Strikingly, sequences in specific families were poorly predicted by some pipelines. For example, sequences from the *Pararnavirae* kingdom or *Abyssoviridae* family were challenging for most teams.

The analysis of viruses with DNA-based genomes (Figure 5) revealed a single family, *Finnlakeviridae*, which was challenging for all pipelines. Notably, the VMR dataset included a single sequence associated with that family (i.e., that for Flavobacterium phage FLiP), whereas ten were present in the Expert dataset. Sequences associated with *Microviridae* were also challenging to assign, as well as those of several floating families (e.g., *Fuselloviridae* and *Polydnaviriformidae*). A cause of this poor performance may be hidden diversity within the family. For example, the ICTV currently recognizes only two *Microviridae* subfamilies (*Bullavirinae* and *Gokushovirinae*), falling short of the extensive diversity within the family (dos Santos et al., 2025).

### Classification of viruses with unknown taxonomic position

Besides the VMR and Expert datasets for which taxonomic information was available, we included 3,151 predicted viral contigs with unknown taxonomic affiliation, all found in shotgun metagenomes in the MGnify database (Figure 1D). To assess how consistently these viruses were assigned, we compared the annotations by different tools and contrasted their Shannon entropy values with those of the VMR dataset (Figure 6AB). The assignment entropy was lower for sequences in the VMR dataset, illustrating that the different pipelines generally agreed on the assignment of sequences with a known taxonomy, but less so for unknown ones that were missing from the training or reference databases.

**Figure 6:**
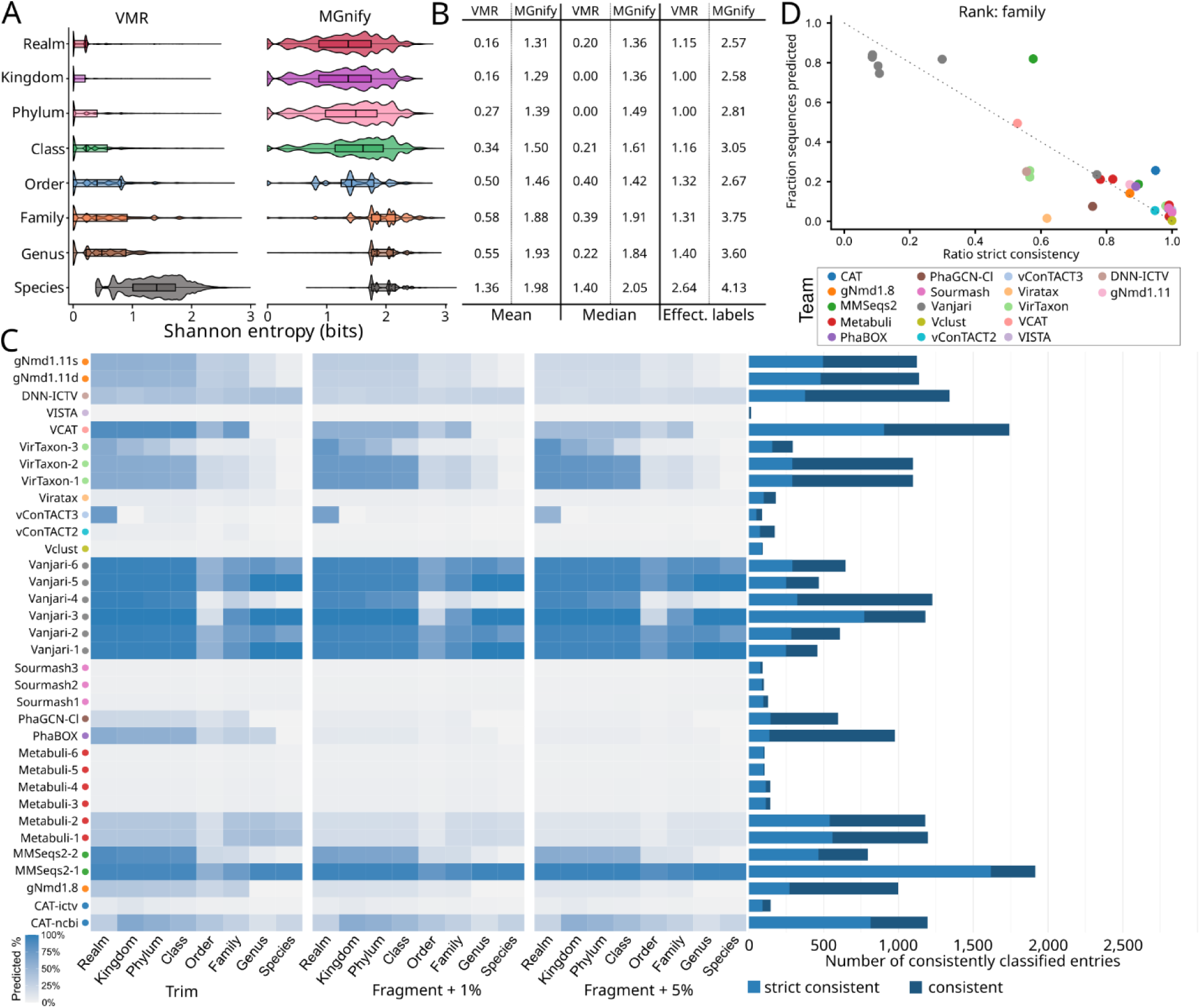
(A) Distributions of Shannon entropy scores for assignments of sequences in the VMR and MGnify datasets at different taxonomic ranks. **(B)** Mean and median Shannon entropy scores and estimated number of effective labels (median) at each taxonomic rank. **(C, D)**: Three sequences were derived from each of the 3,151 initial MGnify contigs, and underwent the operations “trim”, “fragment +1% substitutions, or “fragment +5% substitutions” (Figure 1). When the classification was the same for three operations, or for two operations and a blank for the third, it was defined as strict consistency. When assignment is present for a single operation and blank for the other two, it was defined simply as consistent. **(C)** Heatmap showing the percentage of sequences assigned by each pipeline in each category; bar chart shows the number of consistently assigned entries for the family rank. Strictly consistent in light blue (at least two out of three sequences being assigned consistently, the third can be blank), and simply consistent in dark blue (one assignment and the two others are blank). **(D)** Fraction of assigned sequences versus ratio strict consistency (the number of perfect agreement for the three sequences when at least two sequences out of three have been predicted). The graph shows this at the family rank (see Supplementary Figure 4 for other ranks).

We next assessed the assignment consistency among the three sequences derived from each MGnify contig, to which different transformation operations were applied (“trim”, “fragment + 1% substitutions”, “fragment + 5% substitutions”, see Figure 1). For each pipeline, we calculated the fraction of cases for which assignments were consistent and defined strict consistency as at least two out of three sequences being assigned consistently (the third can be blank). When only one sequence was assigned and the other two were assigned as blank, we called it simply consistent (Figure 6C). Only few MGnify contigs were classified by most tools (Figure 6C), especially in comparison with the VMR dataset (See Figures 4 and 5 at family rank). Exceptions were Vanjari and MMSeqs2-1, which assigned most of the MGnify contigs, albeit less consistently than pipelines with lower assignment rates (Figure 6D, Supplementary Figure 3). Vanjari assigned the most MGnify contigs but also had the greatest assignment inconsistency amongst the three connected segments.

Next, we selected which assignments of the novel viral sequences might be most reliable. Figures 4D and 5D show how many of the MGnify sequences were assigned to each family with high confidence [accuracy * (1 - FDR) > 0.9]. These results were integrated into an ensemble prediction that combines all family-specific accuracy scores and FDR metrics (Figure 7A). In total, 498 sequences from 148 families were retrieved from the “trim” section of the MGnify dataset (*n*=3,151). To benchmark the assignments with a tree-based approach and prior to adding MGnify sequences, we generated viral proteomic trees (ViPTrees) for the sequences that were assigned within each realm by using genome-wide sequence-similarities based on tBLASTx (Nishimura et al., 2017). This enabled assessing whether the sequences assigned to families within the realms formed monophyletic clades. From the VMR dataset, we randomly selected five sequences (or all available if fewer) for each of the families and realms and built one tree per realm (Supplementary Figure 5). This information was used to filter families for which monophylies could not be verified by this method. In a second step, the MGnify sequences within the families predicted through the ensemble method were selected and the tree was rebuilt, this time including both selected reference and these ensemble sequences (Figure 7B, Supplementary Figure 6). The monophyletic clades within the *Monodnaviria* and *Adnaviria* realms are shown in Figure 7B (see Supplementary Figure 6 for all realms). The analysis (Figure 7A) revealed that nine and four families contained MGnify sequences obtained using the ensemble method, yielding six and three monophyletic clade assignments, respectively. Considering all realms together (Figure 7C, Supplementary Figure 6), monophylies confirmed the classification of 212/498 sequences (42.6%) selected in the ensemble MGnify pipeline.

**Figure 7:**
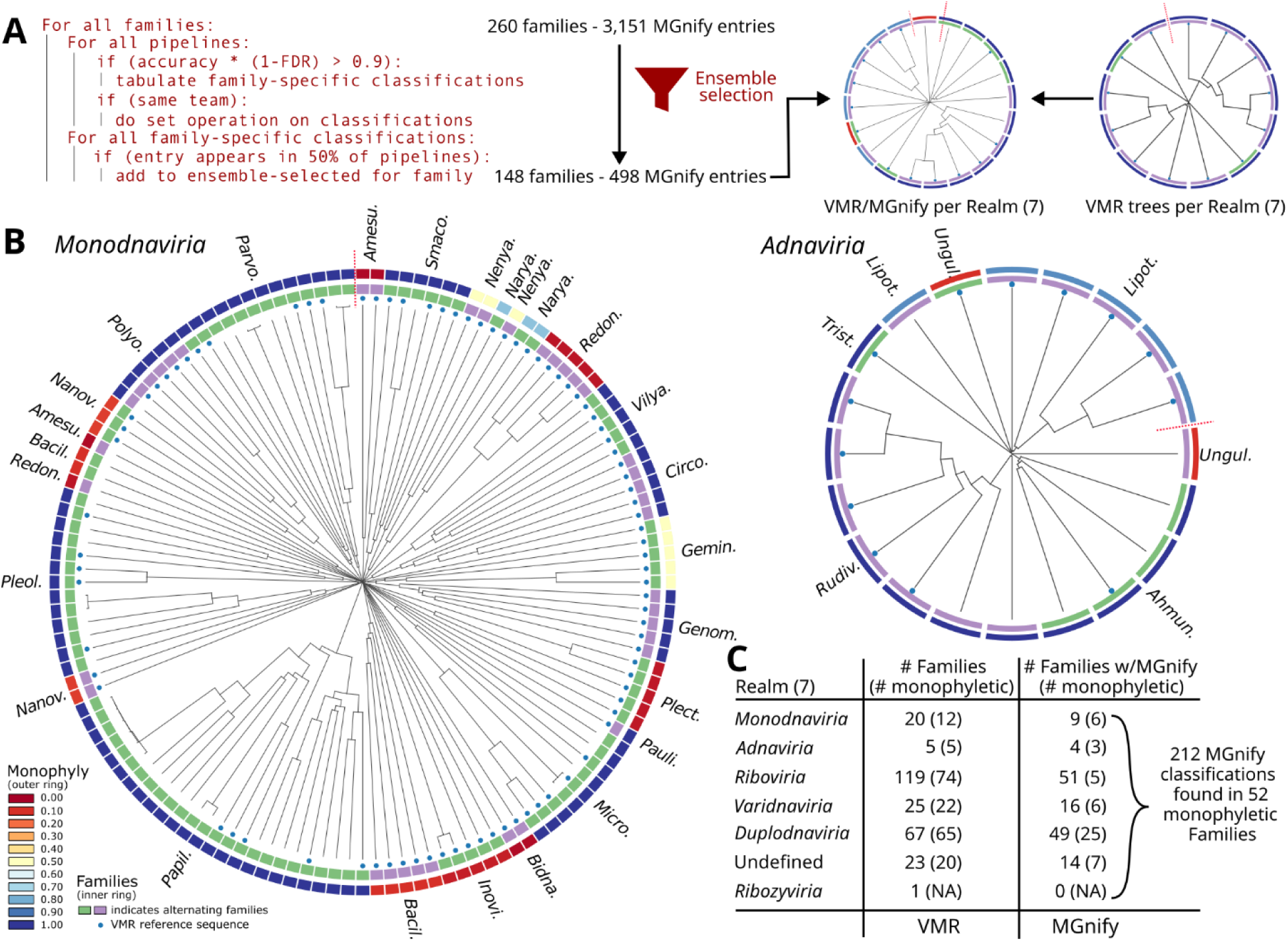
(A) Pseudo-code of the ensemble selection (left-hand side) and validation pipeline with VipTree (right-hand side). **(B)** Circular ViPTrees for the Monodnaviria and Adnaviria realms (for other realms, see Supplementary Figure 6), including “trim-only” sequences from the MGnify dataset. These sequences were assigned to families in the realms via the ensemble selection process and (up to) five randomly selected VMR reference sequences per family (blue dots). The outer ring represents the monophyly ratio of each family, i.e., the ratio between the number of leaves for a given family divided by the number of leaves within the clade of the Most Recent Common Ancestor (MRCA) that includes all members of that family. The inner ring alternates colors to highlight different families. **(C)** Summary of the number of families per realm for the VMR-only and MGnify+VMR data. Only families with sequences assigned by the pipelines are indicated (monophyletic families between parentheses).

### Usability and computational performance

An important goal of the ICTV Computational Virus Taxonomy Challenge was to bring together reproducible virus taxonomy pipelines that are freely accessible in the public domain. Thus, all scripts, pipelines, and configuration files are available through GitHub (Table 1, Supplementary Table 5). To assess the openness, reproducibility, and usability of these resources, a ten-point evaluation framework was applied, encompassing repository accessibility, documentation quality, installation guidance, dependency management, automation, issue tracking, and overall organization. All repositories were accessible and included clear documentation with installation instructions, enabling deployment in both local and high-performance computing (HPC) environments. While the degree of automation and workflow integration varied, many repositories were well-structured and implemented best bioinformatics practices that support reproducible and transparent software use.

To benchmark the different pipelines, we accessed all submitted GitHub repositories and tested them on an HPC cluster, using the same CPU or GPU node for consistency. We observed substantial differences in runtime and memory requirements among the pipelines (Figure 8, Supplementary Table 5). For example, Sourmash, Vclust, and Viratax were very fast and efficient (1.5, <5, and <6 minutes, with 1.5 GB, <16 GB, and <0.2 GB RAM, respectively). Metabuli variants were also very fast (3–6 minutes) but with high memory requirement (>70 GB RAM), although we note that the maximum memory can be set with the “--max-ram” parameter, enabling it to run on consumer grade computers (Lee et al., 2025). The performance of Vanjari depended strongly on model configurations, with its ensemble and fast models taking about one hour but with high memory usage (≈76 GB RAM), whereas its main models failed to complete within 72 h despite relatively low memory consumption (≈16 GB RAM). VISTA also did not complete within the 72-h time limit (≈52 GB RAM).

**Figure 8:**
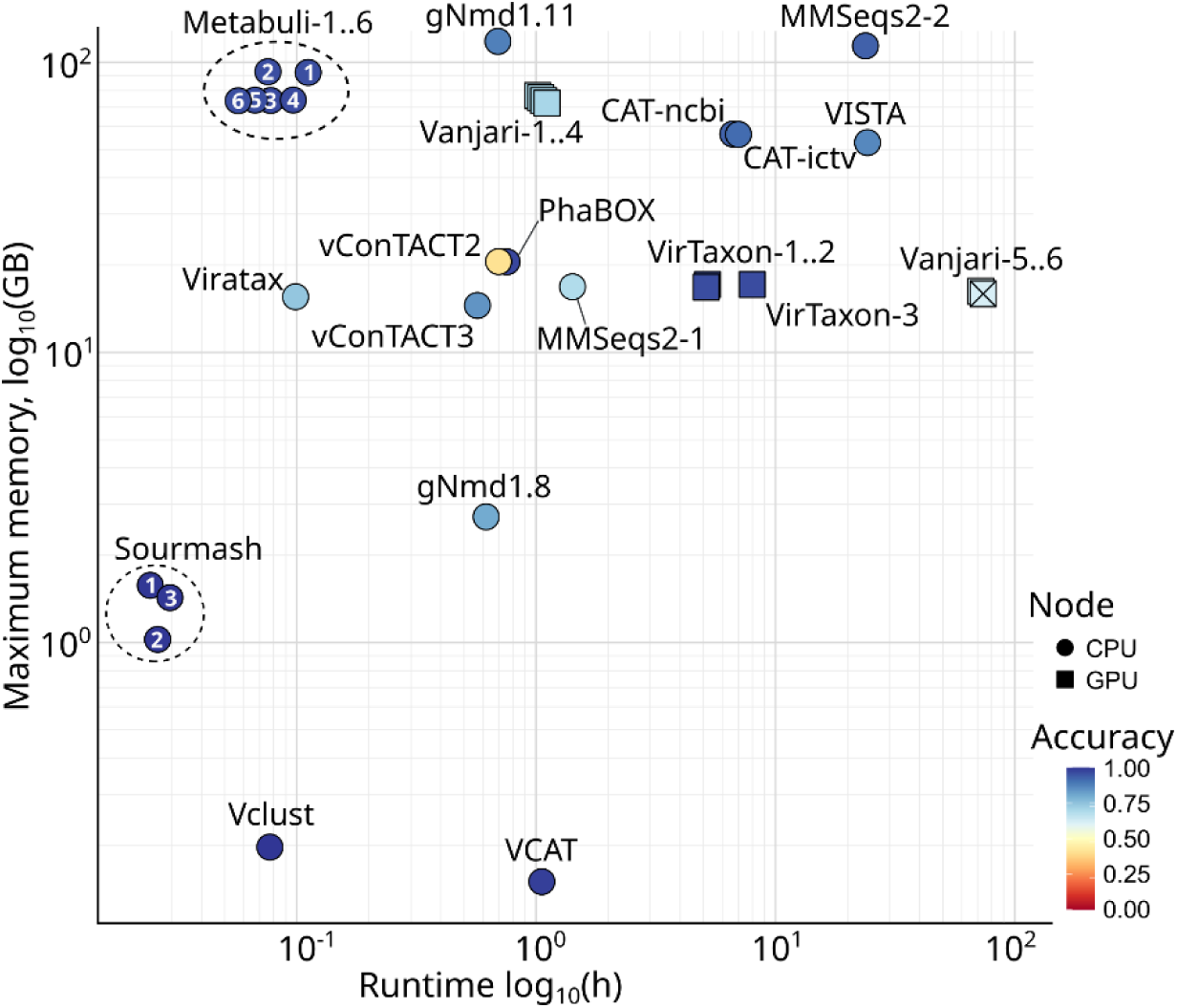
Runtime and peak memory usage of all entries to the ICTV Taxonomy Challenge. Pipelines were downloaded from the respective GitHub repositories and ran on the 59,907 sequences in the challenge dataset, either on a CPU node (40 CPUs, 100 GB RAM) or a GPU node (40 CPUs, 100 GB RAM, NVIDIA H100) on a high-performance computing cluster. Jobs that exceeded three days of wall-clock time were terminated and are not shown crossed on the graph (Vista, Vanjari5-6), as the runtimes were capped at 72 h. The accuracy displayed was calculated for the predicted fraction of the VMR dataset at the family rank (Figure 2B).

Overall, the computational benchmarking revealed three broad categories: (i) fast pipelines with high memory requirements (Metabuli), (ii) moderate runtime pipelines with moderate to high memory requirements (CAT, PhaBOX, vConTACT2, and VirTaxon), and (iii) fast and memory-efficient pipelines (geNomad, Sourmash, Vclust, VCAT, Viratax, and MMSeqs2-1). In addition, some pipelines had clear limitations: VISTA and the main Vanjari versions ran prohibitively slowly, whereas MMSeqs2-2 and geNomad 1.11 had high memory requirements. These trade-offs underscore that pipeline selection will also need to consider the availability of computational resources and time constraints.

## Discussion

### Classifying the virosphere

This first ICTV Computational Virus Taxonomy Challenge asked an important question for modern virology: to what extent can automated, high- throughput computational tools assign unannotated viral sequences to the fifteen ranks in the ICTV virus taxonomy? The dataset was composed of sequences derived from reference genomes of classified viruses (VMR dataset), sequences of previously unclassified viruses (Expert dataset), and hypothetical virus sequences predicted from metagenomic data (MGnify contigs). The VMR dataset generally had high assignment rates, suggesting that most tools perform well in assigning known viruses, whereas the Expert and MGnify datasets proved to be more challenging. Together, the dataset enabled a rigorous benchmarking of the strengths and limitations of current bioinformatic approaches.

The submitted pipelines varied greatly on the evaluated metrics of accuracy, false discovery rate, and computational performance (time/memory). The ability of fast tools with low or configurable memory requirements, such as Vclust, Sourmash, Metabuli, and VCAT, to accurately assign known virus sequences, even when fragmented or mutated (VMR dataset), suggests immediate utility in clinical and environmental surveillance pipelines or rapid detection of known pathogens. Pipelines such as geNomad, MMSeqs2-2, Metabuli, PhaBOX, PhaGCN Cluster, vConTACT2, and VCAT maintained accuracy for the Expert dataset, albeit with variable sensitivity, supporting their use for characterizing sequences from unexplored viromes. As these viruses await official classification at lower taxonomic ranks, the sequences from the Expert dataset were more accurately assigned at higher ranks starting at the family rank. VirTaxon uses a hybrid approach that integrates alignment-based and deep-learning methods, resulting in high accuracy at the family and order ranks within the Expert dataset.

The assignment gap between known and novel viruses remains substantial. While a significant fraction of unknown sequences from the MGnify dataset could be consistently placed into established families, many also remained unassigned. These sequences may represent deep-branching taxa whose affiliation is unresolved by the current ICTV framework. Also, the sequences in the MGnify dataset were predicted to be virus-derived, but they may not be. In metagenomic datasets, a common source of misassignment consists of human, bacterial, and other host-derived sequences, which may be spuriously interpreted as viral. An important caveat in the design of this first Virus Taxonomy Challenge was that we assumed that all sequences were derived from viruses, which made it possible for tools such as VCAT to utilize a small, targeted reference database consisting only of sequences belonging to ICTV-classified viruses. When a full database is used, contaminating sequences from hosts or other cellular organisms may be more readily detected, but it also complicates the classification of virus sequences, which may be spuriously annotated. This is illustrated by the CAT-ncbi results, since this pipeline used the full NCBI non-redundant protein database and did not attempt to filter out non-viral hits.

### Technical considerations

More often than expected, pipeline results included illegal labels (e.g., “Flaviviridae-like”, “Cluster 19”) and invalid combinations of labels along the taxonomic lineage (e.g., a genus assigned to the wrong family). Such anomalies have a direct negative impact on accuracy but are easy to prevent, as they can be automatically detected. Another limitation that became apparent during the evaluation of the pipelines is related to the progressing nature of virus taxonomy, and its yearly assessment by experts. Taxon names at different ranks may change, e.g., they may be abolished (removed entirely), renamed (given a new name), demoted (moved to a lower rank), promoted (moved to a higher rank), split into two or more taxa, or merged (Simmonds et al., 2025). Ideally, a classification pipeline would allow users to specify which MSL version to use, e.g., by implementing a command line parameter such as “--msl_version 39v4” or “--msl_version latest”, which would make it more sustainably applicable. Currently, VCAT is the only pipeline that enables users to choose between MLS39 or MSL40 with the “--dbversion” option. Automatically updating the reference database would benefit greatly from a single access point for downloading all ICTV-classified sequences (i.e., those listed in the VMR or MSL), which is possible in the recently published VirJenDB database (Saghaei et al., 2026).

A critical challenge for automated virus classification is the inconsistency, and sometimes even the absence, of species and genus demarcation criteria, even for ICTV-ratified taxa (Box 1AB). One of the premises of this Taxonomy Challenge was that viruses are so diverse that no single taxonomic method can be used to classify them all (Simmonds et al., 2023). However, only few of the pipelines were biased for specific families or realms, even though they used very different computational methodologies (Extended Methods). This holds promise for computational virus taxonomic classification and suggests that diverse virus sequences may be confidently assigned with a single approach. Some ICTV Subcommittees and Expert Groups have already adopted genome-based criteria that standardize demarcation of low-rank taxa, supported by evolutionary analyses (Chiumenti et al., 2021; Deng et al., 2022; Turner et al., 2021; Van Doorslaer, 2022). As the virus taxonomy remains a living project that depends on the contributions of experts around the world who feed it with new insights, we hope that the findings described above, including any inconsistencies between the pipeline assignments and the ICTV taxonomy, will inspire these experts to dig deeper and unravel the underlying causes and patterns, further improving the taxonomy.

### Outlook

In the ICTV Taxonomy Challenge, the focus was to assess the assignment of unlabeled virus sequences into the established ICTV taxonomy using computational methods. Future studies might also assess the potential of computational tools to suggest novel taxa at all ranks, or even the need for additional ranks. An important direction for future computational research is thus to shift from assigning sequences to established taxa to proposing and delineating novel taxa. Integration of reproducible, data-driven computational methods is essential to transition the virus taxonomy into a sustainable platform that can scale with the rate of virus discovery. As the availability of complete genome sequences is a main criterion for approval of a taxonomy proposal by the ICTV, completeness evaluation would be a critical aspect of such tools.

An important finding of our benchmarking study is the observation that for most tools, viruses were more accurately classified in large families with many reference sequences than in smaller families. This highlights the importance of further charting the virosphere, and the value for ICTV of validating and formally classifying rare viruses associated with less well-described taxa. The availability of multiple and diverse genome sequences will clarify the genomic profile of these taxa and improve the overall performance of automated classification tools.

This first ICTV Computational Virus Taxonomy Challenge highlights the rapidly maturing field of virus bioinformatics. As virus discovery accelerates, tools and pipelines with complementary strengths will provide a foundation for scalable, reproducible, and replicable classification of known and novel viruses. Continued community-driven benchmarking and close collaboration between computational developers and taxonomic experts, promises a future where computational methods enable us to keep pace with the expanding view of the virosphere, actively supporting a more complete, accurate, and adaptable virus taxonomy.

## Materials and Methods

### Organization of the community-driven challenge

The foundations for community engagement in the ICTV Computational Virus Taxonomy Challenge were laid during the ICTV Computational Taxonomy workshop that was organized in the Summer of 2023 in Jena, Germany. The workshop gathered virologists and bioinformaticians for two days of discussion about the future of virus taxonomy (https://evbc.uni-jena.de/events/virus-taxonomy-workshop/). Virologists from multiple ICTV Study Groups were contacted and requested to weigh in with their expertise as part of the Challenge. Following this initial interaction, and upon receipt of expert classifications of the novel viral sequences, we developed a website to communicate the challenge, specify its goals, and provide access to the dataset of unlabeled query sequences (https://ictv-vbeg.github.io/ICTV-TaxonomyChallenge/).

### Description of the input datasets used in the challenge

Three separate datasets were used in the challenge: i) VMR dataset: GenBank accession numbers listed in the ICTV Virus Metadata Resource file VMR_MSL38_v2 were used to download the corresponding genome sequences from NCBI GenBank, yielding 12,624 entries (Sayers et al., 2025). Because some genome sequences (e.g., those of segmented viruses) are covered by multiple GenBank files, the final yield of sequences of known taxonomic affiliation was 15,978 nucleotide sequences. We subsequently removed 78 sequences from version MSL39v4 of the ICTV taxonomy that was published by the end of the challenge submission period; ii) Expert dataset: 840 viral sequences were retrieved from a curated subset of the IMG metagenome database (Chen et al., 2023; Fong et al., 2024; Martinez-Garcia et al., 2024). These sequences were tentatively assigned to the following established taxa or groups: *Amalgaviridae*, *Caudoviricetes*, *Cressdnaviricota*, DJR phages, *Finnlakeviridae*, *Fuselloviridae*, *Huolimaviricetes/Laserviricetes, Huolimaviricetes*, *Microviridae*, *Preplasmiviricota*, *Naldaviricetes*, *Parvoviridae*, virophages, and polinton-like viruses. We subsequently contacted ICTV Study Group experts to aid in the taxonomic classification of these sequences, who confirmed the proposed classifications for 62% of this dataset (521 sequences); and iii) MGnify dataset: for this dataset, we downloaded all 16,702 publicly available assembled metagenomes from the MGnify database as of August 2023, predicted contigs of viral origin with Jaeger (total: 16,152,059 contigs), and provided a preliminary taxonomic classification with geNomad (Camargo et al., 2024; Richardson et al., 2023; Wijesekara et al., 2024). The preliminary geNomad classifications were used to obtain a diverse set of contigs between 5 kb and 300 kb: we randomly selected 20 contigs from each classified taxon (at any rank) that was represented by >20 contigs, 10 contigs from taxa represented by more than 10 but less than 20 contigs, all the contigs from the remaining classified taxa, 500 of the contigs that were only classified as “Virus”, and 1,000 contigs that had no geNomad classification. This yielded a total of 3,151 metagenomic contigs.

### Creating the dataset of sequences for the challenge

The combined dataset was duplicated, enabling different operations, and systematically edited to introduce errors typically found in virus contigs discovered in metagenomic assemblies (Figure 1). In half of the data (one duplicate of the full dataset), sequences were split into four equal fragments, of which two were randomly selected, and 1% or 5% single nucleotide substitutions were randomly introduced, respectively. Sequences in the other half of the data were processed depending on the source: i) sequences from the VMR dataset were trimmed, rotated, reverse-complemented, or not modified, ii) sequences from the Expert dataset were not modified, and iii) sequences from the MGnify dataset were reverse-complemented or not modified. The final dataset available to the participants consisted of 59,907 sequences, from which 234 (i.e., 3×78) were subsequently removed as they were abolished in ICTV MSL39v4 used for the challenge. This dataset was provided as fasta files with anonymized entry headers.

### Evaluation of the classification metrics

The scoring of the entries was divided into two parts. We first focused on sequences for which “ground truth” taxonomy information was available, either because the sequences were derived from viruses previously classified by the ICTV (VMR dataset), or classification was proposed by ICTV Study Group experts (Expert dataset). Then we focused on sequences for which taxonomy information was not available (MGnify dataset). Accuracy was calculated at all ranks by tallying exact matches of the annotations with the ground truth dataset. Accuracy was further broken down by the various sequence transformation operations (see previous section) and by dataset (VMR, Expert, or combined). Unavailable taxa of any ranks were encoded as “nan” in the ground truth dataset. When considering the predictions, missing names (empty string) or the value “NA” were considered as equal. We calculated the False Discovery Rate (FDR) by considering explicit calls only, i.e., not considering missing names, with FDR = (false positives) / (true positives + false positives).

### Benchmarking the classification of unknown sequences (MGnify dataset)

To assess the predictions of entries related to the MGnify dataset (9,454 sequences available, see Figure 1), the following benchmarking strategies were applied: 1) calculating the Shannon entropy of the predictions *between-pipelines*, 2) comparing the consistency of predictions *within-pipeline* across the transformation processes (“trim”, “fragment +1% substitutions, and “fragment +5% substitutions”), and 3) an ensemble method to select sequences with consistent predictions across multiple pipelines, followed by comparative genomics with known viral sequences to assess their assignment. For the latter benchmarking strategy, initial VMR-only trees were calculated for each of the realms except *Ribozyviria* (for which no sequences were predicted to belong across all pipelines). This was done by selecting five sequences (or all if fewer were available) at random from the VMR dataset for each of the component families and analyzing them using VipTree v1.1.2 with default options (Nishimura et al., 2017). Resulting BIONJ trees were loaded into R, mid-point-rooted, and visualized together with the metadata using ggtree v3.10.1 (Yu et al., 2017). Monophyly analysis was performed using the ape::is.monophyletic() function in ape v5.8 (Paradis & Schliep, 2019). The monophyly ratio was defined as follows: for a given family (F), take all the related tips in the tree (size(F)), and determine their Most Recent Common Ancestor (MRCA). The monophyly ration is then: size(F)/size(MRCA(F)). In a second step, the sequences from the MGnify dataset that had undergone the “trim” operation were analyzed using an ensemble approach. For each family and for each pipeline, the metric (accuracy * (1 - FDR)) was used to subset predicted sequences, retaining sequences where the metric was greater than 0.9. When a team provided multiple pipelines, a union operation was applied to keep a single representative sequence. The sequences were then kept if they were predicted to belong to a family by >50% of the pipelines and were added to the VMR sequences to build VMR+MGnify trees, which were then assessed again for monophyly.

### Benchmarking computational performance

Computational performance was benchmarked using the full Taxonomy Challenge dataset, consisting of 59,634 sequences in fasta files. Each pipeline was executed either on a CPU node (40 CPUs, 100 GB RAM) or a GPU node (40 CPUs, 100 GB RAM, NVIDIA H100) on a High-Performance Computing (HPC) cluster. Default parameters were used unless a tool-specific configuration was required (for version information, key commands, and other details see Supplementary Table 5). Runtime and peak memory usage were recorded. Jobs that exceeded three days of wall-clock time were terminated, and their runtimes were capped at 72 h. PhaGCN Cluster and DNN-ICTV could not be benchmarked due to conflicting installation requirements (see comments in Supplementary Table 5).

## Data availability statement

The ICTV Taxonomy Challenge Dataset and processing scripts are available through Zenodo at https://doi.org/10.5281/zenodo.20694980

## Acknowledgments

Y.B. National Natural Science Foundation of China (Grant No. 32270019). G.B. was supported by resources provided by the Pawsey Supercomputing Research Centre’s Setonix Supercomputer (https://doi.org/10.48569/18sb-8s43), with funding from the Australian Government and the Government of Western Australia. R.B. was supported [in part] by the National Center for Biotechnology Information of the National Library of Medicine (NLM), National Institutes of Health (NIH). The contributions of the NIH author(s) are considered Works of the United States Government. The findings and conclusions presented in this paper are those of the author(s) and do not necessarily reflect the views of the NIH or the U.S. Department of Health and Human Services. A.P.C. was supported by the São Paulo Research Foundation (FAPESP, grant 2021/10577-0) and the National Institutes of Health (NIH, grant 5U01DE034196-02).L.D.C. was funded by the Research Foundation Flanders (11L1325N). S.D. was supported by the National Science Centre, Poland, project DEC-2022/45/B/ST6/03032. B.E.D. was funded by the European Research Council (ERC) Consolidator grant 865694: DiversiPHI, the Deutsche Forschungsgemeinschaft (DFG, German Research Foundation) under Germany’s Excellence Strategy – EXC 2051 – Project-ID 39071386, the Alexander von Humboldt Foundation in the context of an Alexander von Humboldt-Professorship founded by the German Federal Ministry of Education and Research. R.A.E. was supported by awards from the Australian Research Council DP250103825 and FL250100019. A.G. was supported by the National Science Centre, Poland, project DEC-2022/45/B/ST6/03032. L.K. was funded by the European Union, ViroInf grant number 955974, APPEAL grant number 101137311. J.K. was funded by the National Research Foundation of Korea (grants 2020M3-A9G7-103933, RS-2021-NR061659, RS-2021-NR056571 and RS-2024-00396026), Samsung DS research fund, Creative-Pioneering Researchers Program and AI-Bio Research Grant through Seoul National University, Novo Nordisk Foundation (NNF24SA0092560). C.L. acknowledge a Marie Skłodowska-Curie Actions Postdoctoral Fellowship from the UKRI Horizon Europe Guarantee program (grant agreement no. EP/Y029585/1), Deutsche Forschungsgemeinschaft (DFG, German Research Foundation) under Germany’s Excellence Strategy – EXC 2051 – Project-ID 390713860. E.J.L. was supported by the National Institute of Allergy and Infectious Diseases of the U.S. National Institutes of Health under Award Number U24AI162625. Y.L. was funded by the National Natural Science Foundation of China (Grant No. 32270019). J.R. was funded by the Swedish Research Council Starting Grant ID 2023-03310_VR. L.R. acknowledges the contribution and support from the Italian national Node (MIRRI-IT) of the European Research Infrastructure MIRRI-ERIC. S.S. acknowledges support from the Mississippi Agricultural and Forestry Experiment Station (MAFES), USDA-ARS agreement 58-6066-3-044, USDA-NIFA SCRI Grant 1029242, and NIFA-USDA Hatch Project 7006130. M.S. was funded by the National Research Foundation of Korea (grants 2020M3-A9G7-103933, RS-2021-NR061659, RS-2021-NR056571 and RS-2024-00396026), Samsung DS research fund, Creative-Pioneering Researchers Program and AI-Bio Research Grant through Seoul National University, Novo Nordisk Foundation (NNF24SA0092560). Y.S. High performance computing support from City University of Hong Kong, the Hong Kong Innovation and Technology Fund (ITF) [MRP/071/20X], RGC GRF CityU 11209823, and CityU 9667256. L.T. was supported by the National Natural Science Foundation of China (Grant No. 32270019). R.T. was supported by The University of Melbourne’s Research Computing Services and the Petascale Campus Initiative. A.V. is partly funded by a National Science Foundation grant # 2412446. Y.W. was funded by the European Union, ViroInf grant number 955974, APPEAL grant number 101137311. A.Z. was funded by computational grant number pl0074-02 from the Poznan Supercomputing and Networking Center. S.R. thanks Patrick Chain, Manuel Martinez-Garcia, Thomas Mock, and their respective teams, for agreeing to share specific viral sequences identified in their unpublished datasets as part of this challenge. The work conducted by the U.S. Department of Energy Joint Genome Institute (https://ror.org/04xm1d337), a DOE Office of Science User Facility, is supported by the Office of Science of the U.S. Department of Energy operated under Contract No. DE-AC02-05CH11231.

## Boxes

### Box 1

**Box 1A:**
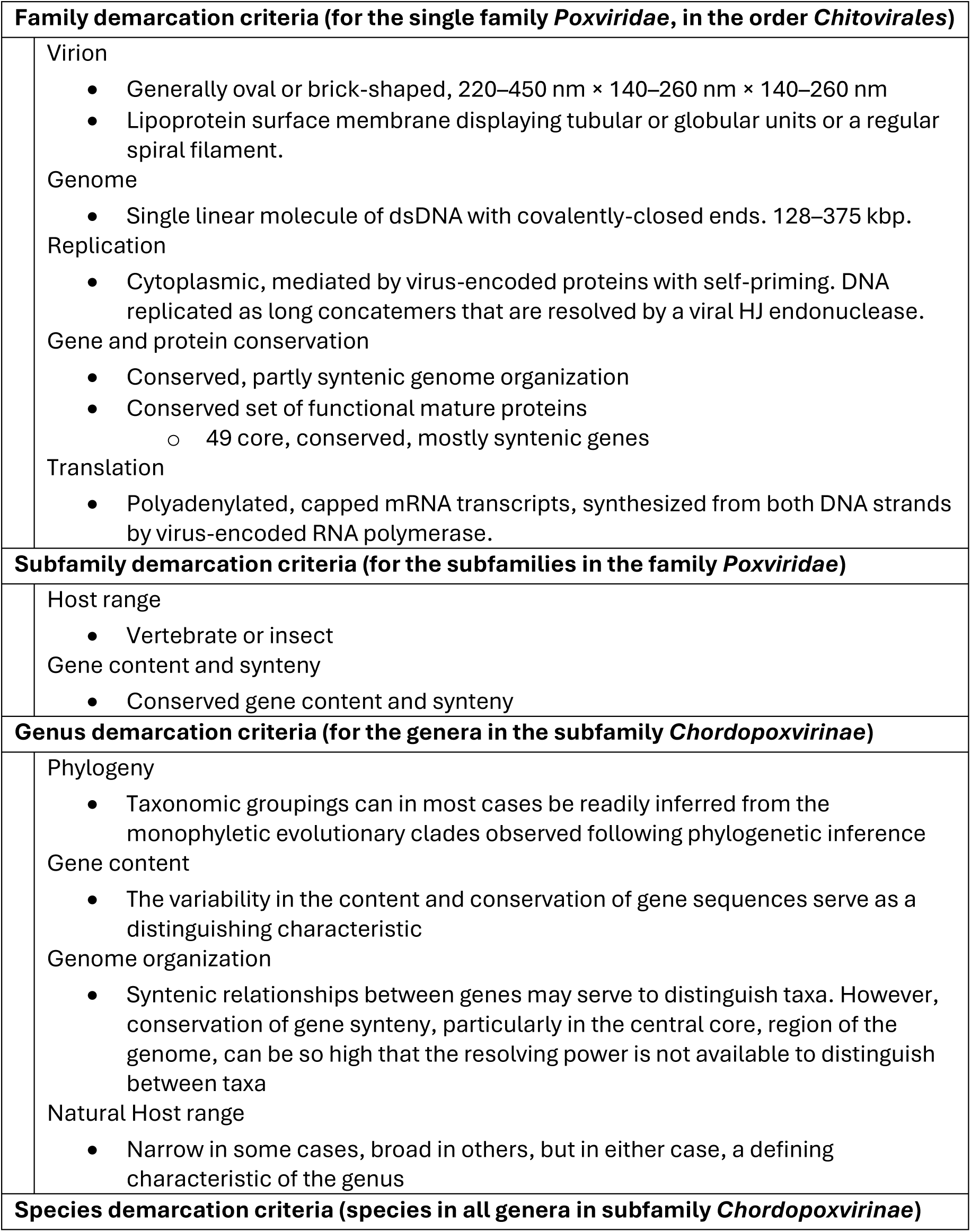
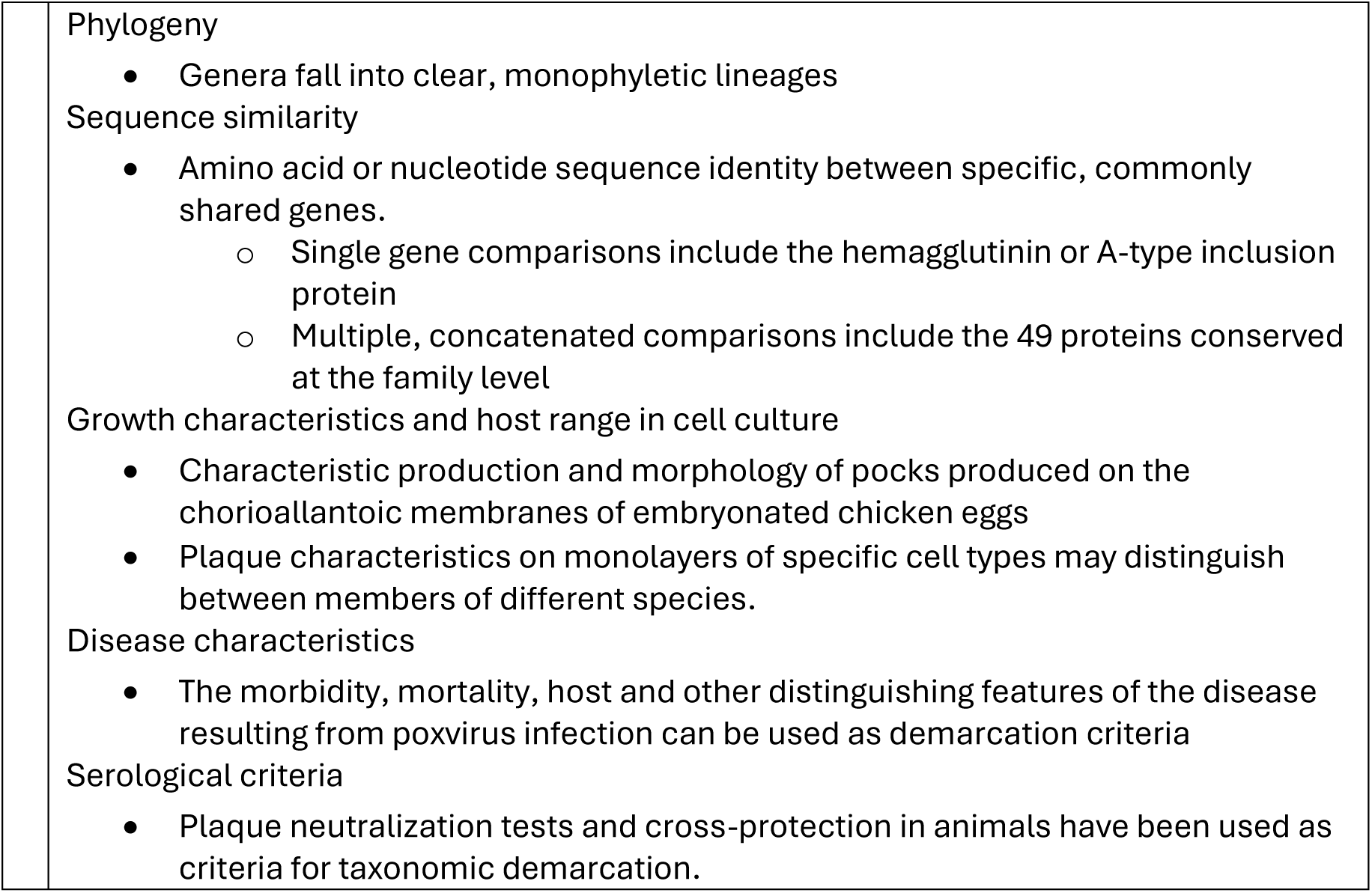
DNA virus order Chitovirales Taxon Demarcation Criteria.

| **Family demarcation criteria (for the single family *Poxviridae*, in the order *Chitovirales*)** |  |
| --- | --- |
| Virion | • Generally oval or brick-shaped, 220–450 nm × 140–260 nm × 140–260 nm<br>• Lipoprotein surface membrane displaying tubular or globular units or a regular spiral filament. |
| Genome | • Single linear molecule of dsDNA with covalently-closed ends. 128–375 kbp. |
| Replication | • Cytoplasmic, mediated by virus-encoded proteins with self-priming. DNA replicated as long concatemers that are resolved by a viral HJ endonuclease. |
| Gene and protein conservation | • Conserved, partly syntenic genome organization<br>• Conserved set of functional mature proteins<br>◦ 49 core, conserved, mostly syntenic genes |
| Translation | • Polyadenylated, capped mRNA transcripts, synthesized from both DNA strands by virus-encoded RNA polymerase. |
| **Subfamily demarcation criteria (for the subfamilies in the family *Poxviridae*)** |  |
| Host range | • Vertebrate or insect |
| Gene content and synteny | • Conserved gene content and synteny |
| **Genus demarcation criteria (for the genera in the subfamily *Chordopoxvirinae*)** |  |
| Phylogeny | • Taxonomic groupings can in most cases be readily inferred from the monophyletic evolutionary clades observed following phylogenetic inference |
| Gene content | • The variability in the content and conservation of gene sequences serve as a distinguishing characteristic |
| Genome organization | • Syntenic relationships between genes may serve to distinguish taxa. However, conservation of gene synteny, particularly in the central core, region of the genome, can be so high that the resolving power is not available to distinguish between taxa |
| Natural Host range | • Narrow in some cases, broad in others, but in either case, a defining characteristic of the genus |
| **Species demarcation criteria (species in all genera in subfamily *Chordopoxvirinae*)** |  |

|  |
| --- |
| Phylogeny |
| • Genera fall into clear, monophyletic lineages |
| Sequence similarity |
| • Amino acid or nucleotide sequence identity between specific, commonly shared genes.<br>○ Single gene comparisons include the hemagglutinin or A-type inclusion protein<br>○ Multiple, concatenated comparisons include the 49 proteins conserved at the family level |
| Growth characteristics and host range in cell culture |
| • Characteristic production and morphology of pocks produced on the chorioallantoic membranes of embryonated chicken eggs<br>• Plaque characteristics on monolayers of specific cell types may distinguish between members of different species. |
| Disease characteristics |
| • The morbidity, mortality, host and other distinguishing features of the disease resulting from poxvirus infection can be used as demarcation criteria |
| Serological criteria |
| • Plaque neutralization tests and cross-protection in animals have been used as criteria for taxonomic demarcation. |

**Box 1B:** RNA virus, order Picornavirales Taxon Demarcation Criteria.

**Box 1B:** RNA virus, order Picornavirales Taxon Demarcation Criteria
| Family demarcation criteria (for the family *Picornaviridae* in the order *Picornavirales*) |  |
| --- | --- |
| Virion | • Non-enveloped, 30–32 nm virions comprising 60 protomers<br>• Capsid proteins composed of three distantly related jelly-roll domains forming icosahedral particles with pseudo-T=3 symmetry |
| Genome | • One positive-sense, monocistronic, single stranded genome segment with a poly(A) tail<br>• Length: 6.7–10.1 kb |
| Gene and protein conservation | • The orthologous proteins 1B, 1C, 1D, 2C, 3C and 3D are conserved between genera<br>• Proteins 1A, 2A, 2B, 3A and 3B are highly divergent among members of different genera.<br>• Conserved genome organization |
| Proteins | • Proteins are translated directly from genomic RNA using an internal ribosomal entry site (IRES)<br>• Primary polyprotein translation product proteolytically cleaved into mature proteins by one or more virus-encoded proteinases |
| Host Range | • Vertebrates |

| **Subfamily demarcation criteria (for the subfamilies in the family, *Picornaviridae*)** |  |
| --- | --- |
|  | Phylogeny<br>Phylogenetic reconstruction of conserved proteins form monophyletic subfamily evolutionary lineages<br>Genome conservation<br>GRAViTy analysis (Mayne et al., 2024) distinguishes between subfamilies based on phylogenetic analysis and metrics of sequence conservation |
| **Genus demarcation criteria (for the genera in the subfamily *Ensavirinae*)** |  |
|  | Phylogeny<br>Monophyletic phylogenetic lineages<br>Protein<br>At least 50% amino acid identity over the length of the polyprotein<br>VPg protein sequence conservation<br>May possess a Type-I, II, or IV internal ribosomal entry site (IRES)<br>Host<br>Virus replication primarily in (but not limited to) the gastrointestinal tract |
| **Species demarcation criteria (for species in the genus *Enterovirus*)** |  |
|  | Phylogeny<br>Monophyletic phylogenetic lineage<br>Genes<br>Essentially identical genome map<br>Proteins<br>Less than 30% divergent in polyprotein aa sequence<br>Less than 40% divergent in P1 aa sequence<br>Less than 30% divergent in aa sequence of the non-structural proteins 2C + 3CD<br>Share a common genome organization |

