## Supplementary methods for "First community challenge for automated virus taxonomy"

Affiliations: see section at the end of the supplementary methods

### Supplementary Methods

The seventeen pipeline descriptions are organized in the same order as the list of entries received for the ICTV Taxonomy Challenge (Table 1, main manuscript).

#### 1. Methods for CAT-ncbi & CAT-ictv

**GitHub**: https://github.com/MGXlab/ICTV_TaxonomyChallenge_CAT/

For a detailed description of the CAT (Contig Annotation Tool) algorithm, please refer to von Meijenfeldt et al. (von Meijenfeldt et al., 2019).

**Database generation**

CAT automatically downloads the database, in this case the full NCBI non-redundant protein database (nr) (NCBI Resource Coordinators, 2013) and prepares a DIAMOND database (Buchfink et al., 2021). For each protein in nr, CAT finds the last common ancestor of all taxa associated with that protein (e.g., when a protein is encoded by *Escherichia coli* and *Escherichia hermanii*, the last common ancestor will be *Escherichia*). If a protein is not associated with a taxon, it is ignored.

**Taxonomic assignment**

For taxonomic classification, CAT predicts open reading frames on all contigs in the input file using Prodigal. It then performs a protein search against the DIAMOND database, in which all matches are included that have a bit-score within 10% of the best hit. CAT assigns the taxonomy to each ORF by finding the last common ancestor of all organisms with top hits. It then assigns taxonomy to a contig by adding up the bit-score evidence for each taxon found across all ORFs and reports it if the bit-score evidence exceeds a threshold. For the ICTV Taxonomy Challenge, we ran CAT 6.0.1, which uses Prodigal 2.6.3 (Hyatt et al., 2010) and DIAMOND 2.1.10 (Buchfink et al., 2021). We queried the nr database of December 12$^{th}$, 2024. In the CAT-ncbi entry, the CAT output was used directly, while for the CAT-ictv entry, the ICTV taxonomy was reconciled with ICTV VMR39v4. To do this, we retrieved the NCBI taxids corresponding to the genome sequences whose accessions serve as reference sequences in the ICTV taxonomy. We then made the taxids synonymous with the viral annotation of their respective taxon according to VMR39v4. When multiple taxa were annotated to the same genome, we assigned the last common ancestor.

#### 2. Methods for geNomad1.8

**GitHub:** https://github.com/Janeeeyeo/ICTV_challenge

We used geNomad v1.8.0 (Camargo et al., 2024) with database v1.3 without any customization and according to the geNomad documentation found here: https://github.com/apcamargo/genomad

geNomad is a tool for identifying and classifying viral and mobile genetic element sequences, combining marker gene searches with a neural network to assign ICTV-compatible taxonomy. For the ICTV Taxonomy Challenge, we ran geNomad v1.8.0 using the 'end-to-end' module, which integrates sequence identification and taxonomic classification into a single pipeline. The geNomad database v1.3 was used as the reference for taxonomic assignments. The full challenge dataset was provided as a combined FASTA file and processed with the '--splits 8' parameter, which divides the input into eight partitions for parallel processing, and the '–cleanup' flag, which removes intermediate files upon completion to reduce disk usage. All other parameters were kept at their default values. The taxonomic output from geNomad was used directly for submission without additional filtering or post-processing steps.

#### 3. Methods for MMseqs2-1 & MMseqs2-2

These two contributions both used MMseqs2 (Steinegger & Söding, 2017) to perform sequence searches. The two pipelines differ in the creation of the reference database, search and prefiltering commands, and taxonomy assignment based on the matches.

**GitHub (**MMSeqs2-1): https://github.com/UriNeri/ictv-mmseqs2-protein-database

**GitHub** (MMSeqs2-2): https://github.com/apcamargo/ictv-taxonomy-challenge-nr

**Database generation**

MMseqs2-1: viral protein sequences were downloaded from RefSeq (Goldfarb et al., 2025) and GenBank (Sayers et al., 2024) using their assembly summary tables from NCBI

Genomes (available through https://ftp.ncbi.nlm.nih.gov/genomes/ASSEMBLY_REPORTS/ and accessed on 27/01/2025). Downloaded protein sequences unclassified by the ICTV (i.e., not present in VMR_MSL39v4) were subsequently removed from the database. This amounted to a total of 5,900,899 protein sequences in the database of submission 1 (MMSeqs2-1), hereafter called the 'GenBank database'.

MMseqs2-2: all NCBI nr protein (NCBI Resource Coordinators, 2013) sequences were retrieved from the NCBI nr BLAST database and exported to the FASTA format on February 8$^{th}$, 2025. NCBI taxonomic assignments were then converted to ICTV lineages using taxopy (https://apcamargo.github.io/taxopy/) with a VMR_MSL39v4 taxdump generated with TaxonKit (Shen & Ren, 2021). Each sequence was mapped to an ICTV lineage by identifying the lowest-ranking taxon shared between its NCBI lineage and the VMR_MSL39v4 taxonomy, using that taxon's position within VMR_MSL39v4 to define its ICTV classification. The final 11,301,117 protein sequences were stored in an MMseqs2 database, hereafter called the 'NR database'.

**Taxonomic assignment**

The ICTV taxonomy challenge dataset was searched against GenBank with MMseqs2 taxonomy with a sensitivity of 7 and a maximal e-value of 1e-3. The confidence score was taken from the MMseqs2 output and represented the fraction of protein fragments for that sequence in agreement with the assigned lowest common ancestor (LCA) by the taxonomy command.

In the MMSeqs2-2 approach, ORFs were extracted and searched against the NR database using MMseqs2, with ORF-level taxonomies assigned via the 2bLCA algorithm. These ORF-level taxonomies were then aggregated into genome-level taxonomies using the majority vote approach implemented in taxopy, with votes weighted by -ln(E-value) and capped at a maximum weight of 1000. Species-level assignments additionally required a minimum average identity of 80% between query and target ORFs. For exact MMseqs2 commands and the genome-level assignment script, see https://github.com/apcamargo/ictv-taxonomy-challenge-nr/blob/master/scripts/mmseqs2_search.sh

The final taxonomy of the challenge sequences was ultimately assigned based on a weighted majority vote implemented in taxopy (https://github.com/apcamargo/taxopy) for all ORFs from the same challenge sequence. The weights for the LCA assignment corresponded to the negative log of the E-value of the alignment. The confidence score of each taxon was calculated as the sum of the weights of the ORFs that were assigned to it, divided by the sum of the weights of all ORFs for that sequence.

**Workflow orchestrations**

MMseqs2-1: concentrated the database generation, search, and result formatting in a single Jupyter Notebook (Kluyver et al., 2016). The evaluated runtime in this benchmark only pertained to the mmseqs taxonomy call.

MMseqs2-2: utilized pixi tasks (Fischer et al., 2025) to download the precompiled database, launch the different MMseqs calls, and execute a custom python script and result formatting. The evaluated runtime in this benchmark pertained to the entire pipeline (`pixi run pipeline`).

#### 4. Methods for Metabuli1 to Metabuli6

**GitHub**: https://github.com/jaebeom-kim/Metabuli-ICTV-challenge

Metabuli is a *k*-mer-based taxonomic classifier (Kim & Steinegger, 2024). Its novel *k*-mer structure, the metamer, jointly encodes amino acid (AA) and DNA information to integrate AA-based sensitive detection of potentially novel viruses with DNA-based precise differentiation of viruses of closely related taxa. Metamers from query and reference sequences are compared to identify exact AA-level matches for sensitive search, while the DNA-level identities of these matches enable specific classifications. Metabuli reports identity scores based on the number of AA matches, DNA Hamming distances, and sequence lengths. We used these scores as confidence scores in this challenge.

Regarding usability, Metabuli supports Windows, macOS, and Linux with a configurable RAM limit (default: 128 GB). It runs efficiently even on an 8 GB RAM notebook with query and database files ≈100x larger than this challenge (Kim & Steinegger, 2024). Additionally, the stand-alone Metabuli App (Lee et al., 2025) provides a GUI for database curation, taxonomic classification, interactive result visualization (Sankey/Krona), raw-read quality control, and clade-specific read extraction.

**Database generation**

NCBI-style taxonomy dump files were created from the ICTV file (available at https://ictv.global/vmr) VMR_MSL39.v4_20241106.xlsx using TaxonKit (Shen & Ren, 2021).

In the initial submission, the database was built using GenBank viral assemblies (genomic.fna.gz) filtered against the ICTV VMR (VMR_MSL39.v4_20241106.xlsx). This yielded 17,963 of 19,769 accessions (~91% of species). However, this filtering unintentionally excluded all accessions from several families (*Abyssoviridae*, *Pseudoviridae*, *Belpaoviridae*, *Rhodogtaviriformidae*, *Brachygtaviriformidae*, and *Bartogtaviriformidae*) and most *Metaviridae* sequences (29 of 31).

For the final submission, the database was built using NCBI Batch Entrez (https://www.ncbi.nlm.nih.gov/sites/batchentrez) to directly retrieve all 19,769 accessions listed in the VMR file. This approach successfully downloaded 19,766 sequences into a single FASTA file. Only three accessions (JAEILC010000038, GECV01031551, and AUXO017923253) were omitted, as the NCBI portal reported these accession numbers as invalid.

**Taxonomic assignment**

Six submissions were generated using different --min-score and --min-sp-score (identity score thresholds for general and species-rank classification, respectively) and --tie-ratio (relative score threshold for LCA determination). Three configurations were tested: default, precise-s (--min-score 0.15, --min-sp-score 0.5), and precise-l (--min-score 0.07, --min-sp-score 0.3). Each configuration was executed with both the default (--tie-ratio 0.95) and conservative (--tie-ratio 0.9) relative score thresholds.

#### 5. Methods for PhaBOX

**GitHub:** https://github.com/KennthShang/PhaBOX/tree/main/ICTV_challenge

PhaBOX is an integrated web server designed for the comprehensive characterization of viral sequences from metagenomic data, encompassing virus identification, taxonomic classification, lifestyle prediction, and host assignment (Shang et al., 2026). The taxonomy module leverages the latest version of PhaGCN, a semi-supervised learning framework that synergizes nucleotide-level sequence features with gene-sharing network topology (Shang et al., 2021). In this network, DNA sequences are encoded into fixed-dimensional feature vectors using a convolutional neural network initialized with a skip-gram-embedding layer to capture local k-mer dependencies; these vectors constitute nodes within a heterogeneous knowledge graph. Edges are established via protein-level homology, utilizing DIAMOND (Buchfink et al., 2021) and Markov Clustering (MCL) to identify orthologous groups, with connectivity defined by a composite metric of shared protein cluster significance and alignment similarity. A graph convolutional network then aggregates features from labeled reference genomes and unlabeled contigs to propagate taxonomic labels. Furthermore, the algorithm enables the *de novo* detection of novel viral lineages via graph clustering. Sequences lacking genus-rank classification are automatically organized into viral Operational Taxonomic Units based on average amino acid Identity and shared protein profiles. The submission in the main text is generated using default parameters (--len 3000), which will filter out contigs <3,000 nt.

#### 6. Methods for PhaGCN

**GitHub:** https://github.com/xiahaolong/PhaGCN_Cluster/tree/main

PhaGCN_Cluster is a GCN-based model that uses deep learning classifiers to learn species mask features for novel virus taxonomy classification (Xia et al., 2026). We used phaGCN_cluster with standard settings and further processing of the results is documented here: https://github.com/xiahaolong/PhaGCN_Cluster/tree/main/results

#### 7. Methods for Sourmash1 to Sourmash3

**GitHub:** https://github.com/sourmash-bio/2024-ictv-challenge-sourmash

Sourmash is a k-mer based multi-tool for sequence comparison (Irber et al., 2024). It relies on the FracMinHash sketching technique to randomly subsample k-mers for fast and lightweight analyses. The default sourmash parameters (DNA k-mer length 31, scaled=1,000) were selected for microbial queries and are not ideal for viruses that have shorter and more diverse genomes. To increase resolution for shorter viral sequences, we increased the FracMinHash sketch resolution by using scaled=50. To increase sensitivity across nucleotide substitutions, in addition to standard DNA k-mers of length 21 (sourmash-2) and 31 sourmash-3), we used an alternative k-mer type: “skip-mers” or “cyclic q-grams”, k-mers built with regular interval gaps (Clavijo et al., 2017). We used skip-mers of length 24 (sourmash-1), built by enabling a gap at every third base, enabling nucleotide mismatches at these positions. As such, a 24 length skip-mer represented an original sequence length of 32 nt, which retains search specificity while increasing sensitivity by allowing mismatches at the gapped positions. We followed a triplet pattern as in Clavijo et al. to tolerate third base pair wobbles when the skip-mer was in the correct frame.

For taxonomic classification of viral sequences, we use the approach outlined in (Portik et al., 2022). First, *sourmash gather* searches FracMinHash sketches of the query sequences against a database of all ICTV VMR viral genome sequences, here set to require a minimum estimated sequence match of 200 nt. Then, *gather* used a greedy minimum set cover approach to select the smallest set of reference genomes that best matched each query sequence (Irber et al., 2022). Finally, *sourmash taxonomy* was used to apply the taxonomic information from these reference genomes to the query sequence above a set match threshold. Here, a classification threshold of 75% estimated log containment (containment ANI; (Hera et al., 2023)) was used to accept classification at a taxonomic rank. If the threshold was not met, an LCA approach was used with the matched reference genome sequences until there was sufficient % match. If a reference genome match was identified by *gather* but the taxonomic threshold was not met, a match was reported as “below threshold”. If no reference genome was found in the *gather* step (i.e., a sequence matching reference genome(s) did not meet the 200-nt search threshold), no taxonomic assignment was reported for the query.

**Parameter tuning**

The *sourmash gather* approach is a reference-based analysis that relies on matching a sequence to closely related reference genome sequences. As reference databases grow and improve over time, so too should the capacity of this method to assign viral sequences. However, this method performs less well for unknown or novel viruses, as in the “expert” set with only family-rank relatives in the reference database. To utilize sourmash for these, future research could assess the use of other sourmash k-mer types, including amino acid k-mers, degenerative protein alphabets, and additional gapped k-mer types (skipm1n3).

As a primary focus of sourmash is to enable analysis on large virome and metagenome datasets, we selected parameters to optimize speed and memory utilization while still providing resolution for relatively small queries. To improve assignment of the shortest sequences in the challenge dataset (≈500 nt and shorter), the FracMinHash resolution could be increased (i.e. ,scaled=25, 10). Note that scaled=1 is feasible but is no longer “sketching”, as it uses all available k-mers; sourmash is not optimized for this use case. For large-scale datasets without short sequence fragments, we suggest scaled=100 for improved computational performance.

#### 8. Methods for Vanjari1 to Vanjari6

**GitHub**: https://github.com/houndry/vanjari

Vanjari (Virus Assessment using Neural networks for Just-in-time Analysis and Rapid Identification) uses two separate models for classification. For the main model, Vanjari divides the input viral sequence into sections that are 1,000 nt long and feature vectors for each subsequence are calculated using Nucleotide Transformer (NT) model (500M parameter, multi-species), fine-tuned on a dataset of viral genome sequences. These feature vectors are stacked and passed to a neural network that applies two linear layers with PReLU activations to each vector. An attention module computes a weighted sum over the transformed subsequence features, producing a single context vector for the entire input sequence. Predictions are made from this context vector using a hierarchical softmax layer (https://github.com/rbturnbull/hierarchicalsoftmax). The hierarchical softmax assigns probabilities for each of the nodes in the taxonomic tree. These probabilities are used as Vanjari's confidence scores. Greedy prediction proceeds down the taxonomic ranks until the probability drops below a threshold. Our submissions used thresholds of 0.0 (i.e., no threshold) and 0.5. A second neural network model does not use the features from the NT model but instead uses a series of convolutional layers before predicting using the final hierarchical softmax layer. Since this second model does not require the NT model, it is substantially faster. Two submissions were made using this faster model with thresholds of 0.0 and 0.5. Vanjari can ensemble outputs from multiple models by averaging the probabilities at each node in the prediction hierarchy, followed by greedy prediction with a threshold. Our fifth submission used an ensemble of the main model and the fast model, with a thresholds of 0.0 and 0.5. We have trained new Vanjari models that incorporate reverse-complement sequences, and we expect this will improve prediction accuracy on those cases.

#### 9. Methods for Vclust

**GitHub**: https://github.com/aziele/ICTV-TaxonomyChallenge-Vclust

Vclust is an alignment-based tool for calculating Average Nucleotide Identity (ANI) between complete or fragmented viral genome sequences and for clustering genome sequences using ANI thresholds (Zielezinski et al., 2025). Because ANI is informative primarily at the species and genus ranks, Vclust operates only within these two taxonomic ranks. Although not designed for taxonomic assignments, Vclust was used in this ICTV Challenge to identify the closest matching genome in VMR for each virus contig and to transfer its ICTV taxonomy to the query. Specifically, global ANI (gANI) values were computed between query contigs and reference VMR genomes (v39.4), and taxonomic labels were assigned *post hoc* based on the highest gANI match. Classification was performed following sequence identity thresholds recommended by the ICTV Bacterial Viruses Subcommittee: gANI ≥95% for species and gANI ≥70% for genera. Contigs below gANI <70% were excluded to prevent speculative higher-rank classification.

#### 10. Methods for vConTACT2

**GitHub**: https://github.com/Rahlfflab/ICTV_challenge

All viral genome files were merged into a single multi-fasta file. Gene prediction was performed using Prodigal v.2.6.3 (Hyatt et al., 2010). Protein sequences were then analyzed with vConTACT2 v.0.11.3 (Bin Jang et al., 2019) alongside the INPHARED phage database of March 1st, 2024 (Cook et al., 2021). Results were compiled using graphanalyzer.py v.1.6.0 (https://github.com/lazzarigioele/graphanalyzer (Pandolfo et al., 2022).

#### 11. Methods for vConTACT3

**GitHub**: https://github.com/bolduc/ICTVTaxoChallenge_vConTACT3

VConTACT3 (Bolduc et al., 2025) is a machine learning-based tool for scalable, hierarchical virus taxonomy that classifies viral sequences across four taxonomic ranks (genus, subfamily, family and order), covering prokaryotic and eukaryotic viruses across four of the six ICTV-recognized realms at the time of this study (*Duplodnaviria*, *Monodnaviria*, *Adnaviria* and *Varidnaviria*). Unlike reference-only classifiers, vConTACT3 establishes classification thresholds from known sequence space via a statistical framework that can be extended to previously uncharacterized viral taxa, allowing novel taxonomic groups to be automatically created where no reference taxonomy exists.

The pipeline starts with gene prediction using pyrodigal, followed by protein clustering via MMSeqs2 at five amino acid identities (30-70% in 10% increments) to construct protein cluster profiles. Pairwise genome distances are calculated from these profiles, which are then used to build gene-sharing networks, which are filtered to remove weakly supported edges and then partitioned by realm. Agglomerative hierarchical clustering is applied to each connected network component using realm- and host-domain-specific distance cutoffs, selected by benchmarking over 60 million parameter combinations against NCBi RefSeq, assigning ranks from order to genus. Novel taxa lacking reference matches are assigned using a standardized naming nomenclature.

**ICTV Challenge**

The 59,907 individual FASTA files in the challenge dataset were concatenated into a single input prior to analysis. VConTACT3 v3.1.7 was run with reference database v223 (https://doi.org/10.5281/zenodo.10935513) and--db-domain of prokaryotes. Since this flag restricts vConTACT3 to using thresholds optimized for prokaryote-infecting viruses, as the challenge dataset included eukaryotic viruses, those sequences were assigned using prokaryote-optimized parameters, representing a known limitation of this submission. All other parameters were defaults. Output was reformatted to the challenge template using the accompanying output_converter.py script.

For a detailed description of the vConTACT3 algorithm, please refer to the publication (Bolduc et al., 2025).

#### 12. Methods for Viratax

**GitHub**: https://github.com/rcedgar/viratax

Viratax was designed to be a fast, lightweight classifier. It emphasizes speed, simplicity and reproducibility over maximizing classification accuracy. The core idea is to generalize rules of thumb relating sequence identity to taxonomic rank. For example, 90% aa or nt identity is often used as a species threshold. Similarly, 75% aa identity is roughly genus, and 50% aa identity is roughly family. These thresholds are gene- and taxon-specific; e.g. 90% aa identity for species is often applied to RNA virus RdRp while other genes diverge more quickly. To address this, every gene in every reference genome (in practice, every ORF) is aligned against all other reference genes to determine ORF-specific thresholds. ORFs in a query contig are aligned to ORFs from reference genomes, and a consensus is taken over aligned ORFs to determine the classification. A fast nucleotide search is performed first to identify hits which are consistent with a species match, i.e. >90% nt identity with >90% coverage of the contig. The confidence estimate reflects identities of the top hits (higher identity gives higher confidence) and consistency (lower consensus gives lower confidence), it is calculated as the weight of the top taxon divided by the sum of weights of all taxa above the rank threshold.

#### 13. Methods for VirTaxon1 to VirTaxon3

**GitHub**: https://github.com/ZhuLab-Fudan/VirTaxonomer

VirTaxon is a two-stage tool for the taxonomic classification of viral contigs, comprising an initial virus identification stage (optional) followed by a taxonomy assignment stage.

Each phase integrates alignment-based methods with machine-learning classifiers. This hybrid design leverages the complementary strengths of the two approaches: the alignment-based module reliably identifies well-characterized viruses, whereas the learning-based module captures taxon-specific sequence features and exhibits greater robustness to sequence fragmentation and evolutionary divergence.

Specifically, the pipeline consists of two stages. During the virus identification stage, each sequence is first aligned to the VMR39v4 reference genomes. Sequences without alignment or only low-quality alignments are then independently assessed by a finetuned ESM2 classifier (Lin et al., 2023). Sequences that either align to known viral references or are predicted as viral contigs proceed to the taxonomy stage, whereas those failing both criteria are labeled non-viral and excluded from further analysis. In the taxonomic classification stage, labels are assigned when a sequence meets predefined genome-level alignment thresholds against reference genomes. For sequences without confident alignment-based assignments, a machine-learning module with seven ESM2 classifiers is applied, with each classifier corresponding to a taxonomic rank from realm to genus. VirTaxon provides three strategies for integrating these classifiers. In the "bottom-up" strategy, predictions start at the genus rank and move upward, and a rank is reported only when its classifier probability exceeds a preset threshold. This approach resolves most sequences at lower ranks and improves runtime. In the "highest" strategy, all seven classifiers are applied, and the final label corresponds to the rank with the highest predicted probability. Confidence scores are derived from minimap2 alignment quality (Li, 2018) or from the probabilities output by the ESM2 classifiers.

Recent updates to VirTaxon incorporated additional information sources to improve taxonomic resolution. In addition to sequence-level alignment (minimap2) and protein-level machine-learning predictions (ESM2), protein-level alignment methods were added to the decision process.

#### 14. Methods for VCAT

**GitHub**: https://github.com/Yasas1994/vcat

VCAT (Virus Contig Annotation Tool) uses a three-step hierarchical sequence comparison strategy for transferring ICTV taxonomy annotations to viral genomes and genome fragments, using MMseqs2 (Steinegger & Söding, 2017) as the primary database search algorithm. This hierarchical framework assumes that biological sequences retain evolutionary signals at different rates. Lower-rank taxa, therefore, tend to share high nucleotide-level similarity, whereas protein- and protein-family–level similarities become more informative for assigning sequences at higher taxonomic ranks.

The VCAT workflow operates as follows: Query sequences are first compared against a nucleic acid database of ICTV genomes, and total Average Nucleotide Identity (tANI) is estimated between the query and all target sequences sharing identical sequence stretches.  Genomes belonging to the same species are assumed to share tANI scores ≥0.81, and those belonging to the same genus are assumed to share tANI scores between 0.81 and 0.49; If tANI is <0.49, or if no matches are detected at this level, the workflow proceeds to the next step. Open Reading Frames (ORFs) are then extracted from the query and compared against a protein database generated from ICTV viruses. A new metric, taxon Average Amino Acid Identity (txAAI), is computed to quantify amino acid–level similarity between the query and each taxon. If txAAI ≥0.49, the query is assigned to the corresponding family or, when necessary, the nearest higher taxonomic rank above family, and if txAAI >0.3, assignment is made at the order rank. Finally, protein profile comparisons are performed using another metric, taxon Average Profile Identity (txAPI). If txAPI ≥0.3, the sequence is assigned to a class, and if txAPI >0.2, it is assigned to the lowest available primary taxonomic rank above class. These cutoffs were derived empirically by comparing ICTV sequences to themselves and examining the taxon-wise distributions of tANI, txAAI, and txAPI. VCAT also provides a workflow that automatically constructs the required nucleotide, protein and profile databases from the latest release of ICTV Virus Metadata Resource (VMR), enabling straightforward updates as new releases become available.

**ICTV Challenge**

The "vcat preparedb" command was used to build the ICTV MSL VMR 39v4 genome, protein, and protein-family profile databases. VCAT was run with default parameters for the ICTV taxonomy challenge, and a custom Python script formatted the results to match the taxonomy challenge template.

#### 15. Methods for VISTA

**GitHub**: https://github.com/YiyunLiu-Ivy/ICTV-TaxonomyChallenge_VISTA

VISTA (Virus Sequence-based Taxonomy Assignment) is an alignment-free method that reconstructs ICTV taxonomy from viral genome sequences and assigns new genomes to established/new taxa. It converts each complete viral genome sequence into a *k*-mer profile based on physico-chemical property sequences, integrating both k-mer frequency and approximate positional information to represent the genome sequence. A two-step feature selection (chi-squared filter followed by extremely randomized trees) reduces these profiles to the most taxonomically informative k-mers. Pairwise distances between genome sequences are then computed using a panel of 19 distance measures, which are normalized to the interval [0,1]. For each family, VISTA applies kernel density estimation and hierarchical clustering to the within-family distance distribution and uses the Fowlkes–Mallows index to identify optimal genus- and species-rank demarcation thresholds and the best-performing distance measure. During taxonomy assignment, VISTA calculates distances between a query genome and all reference genomes in each supported family, as well as the distances among query genomes. It compares the minimum distance of the query to thresholds, and assigns the query to the corresponding species, genus, or discovering potential new species/genus within that family.

For the ICTV Challenge, VISTA mainly focused on viral families whose thresholds were available (38 families plus class *Caudoviricetes*). All challenge contigs were first searched against NCBI *nt* database using *blastn*. Only contigs whose best BLAST hit was a complete viral genome sequence from a VISTA-supported family/*Caudoviricetes* with at least 80% alignment coverage were retained for further analysis. For each such family/*Caudoviricetes*, contigs were grouped and assigned with the VISTA command-line tool using the pre-computed family-specific models and demarcation thresholds. For every contig, its closest reference genome, the minimum normalized distance, and the rank of the assignment (species, genus, family or higher), were recorded. These outputs were then converted into the Challenge template by filling the 15 ICTV ranks based on the predicted rank and by reporting a confidence score defined as (1 - distance).

#### 16. Methods for DNN-ICTV

**GitHub**: https://github.com/BioZhang0831/ICTVTaxoChallenge_model/

This deep learning model has the primary function to classify a set of unknown viral sequences. The model is based on GP-GCN (Gapped Pattern Graph Convolutional Networks) that utilizes RSCU (Relative Synonymous Codon Usage) and TNF (Tetra-Nucleotide Frequencies) to enhance the taxonomic classification accuracy. The model is trained on annotated viruses dataset published by ICTV https://ictv.global/sites/default/files/VMR/VMR_MSL39.v2_20240920.xlsx.

#### 17. Methods for geNomad 1.11d & geNomad 1.11s

**GitHub**: https://github.com/apcamargo/ictv-taxonomy-challenge-genomad/

This submission utilizes a taxonomic pipeline in which geNomad version 1.11 (Camargo et al., 2024) is used to assign input sequences to ICTV taxa. The process involves gene prediction, alignment of the predicted genes to taxonomically informative protein profiles for gene-level taxonomic assignment, and aggregation of gene-level assignments to derive sequence-level taxonomic classifications.

**Database generation**

A detailed description of the procedure used to build the geNomad database can be found in its manuscript (Camargo et al., 2024). Briefly, geNomad's marker database was constructed from an initial collection of 1,425,477 protein profiles. A specificity score was computed for each profile based on its distribution across chromosomes, plasmids, and viruses, and was then used to select a final set of 227,897 markers that were specific to either one of the three classes. Taxonomy was assigned to viral markers by aligning them to viral proteins from the NCBI nr database that were assigned to ICTV lineages using the taxopy library (see methods for the MMseqs2-2 entry). Each marker was assigned to the consensus taxon among its hits, determined using the majority vote function from taxopy.

**Taxonomic assignment**

To assign a query genome to an ICTV lineage, geNomad first performs gene-level taxonomic assignment and then aggregates these assignments into a genome-level taxonomy. Genes are predicted with pyrodigal-gv (Camargo et al., 2024) and queried against the geNomad marker database using MMseqs2 in a protein-versus-PSSM search. The geNomad 1.11d entry used default search sensitivity (-s 4.2), whereas the geNomad 1.11s entry used increased sensitivity (-s 7.5). For each query protein, the taxon is transferred directly from the best-matching marker. These gene-level assignments are then aggregated into a genome-level taxonomy using the majority vote functionality from taxopy, weighted by alignment bitscore, whereby the genome is assigned to the lowest-ranking taxon that accounts for at least 50% of the total bitscore across all genes.

### Affiliations

Cédric Lood[1,2,*], Swapnil P. Doijad[2], Evelien M. Adriaenssens[3], Yiming Bao (鲍一明)[4,5], Jakub Barylski[6], Ben Bolduc[7,8,9], George Bouras[10,11], J. Rodney Brister[12], C. Titus Brown[13], Antonio Pedro Camargo[14,15], Lander De Coninck[16], Sebastian Deorowicz[17], Robert Edgar[18], Robert A. Edwards[19], Shitao Gong[20], Arthur Gruber[21], Adam Gudys[17], Ernestina Hauptfeld[22], Anneliek ter Horst[13], Tianyang Huang[20], Jingzhe Jiang[23], Lars Kaderali[24], Jaebeom Kim[25,26], Mart Krupovic[27], Jens H. Kuhn[28], Elliot J. Lefkowitz[29], Mathieu Leobold[30], Shuai-Cheng Li[31], Yiyun Liu[4,5], F. A. Bastiaan von Meijenfeldt[32], Uri Neri[14], Judit J. Penzes[33], N. Tessa Pierce-Ward[13], Janina Rahlff[34,35,36], Alejandro Reyes Muñoz[37], Luisa Rubino[38], Sead Sabanadzovic[39,40], Jiayu Shang[41], Peter Simmonds[43], Martin Steinegger[25,26,44,45], Matthew B. Sullivan[7,8,9], Yanni Sun[46], Lili Tian (田莉莉)[4,5], Yi-Gang Tong[47], Robert Turnbull[48], Dann Turner[49], Arvind Varsani[50,51], Ziye Wang[52], Yasas Wijesekara[24], Wytamma Wirth[53,54], Haolong Xia[55], Shuo Yang[31], TzeChing Yeo[34], Jinbei Zhang[46], Xianglilan Zhang[56,57], Shanfeng Zhu[20], Andrzej Zielezinski[58], Simon Roux[14], Bas E. Dutilh[2,22,*]

---

[1] Department of Biology, University of Oxford, Oxford, UK

[2] Institute of Biodiversity, Faculty of Biological Sciences, Cluster of Excellence Balance of the Microverse, Friedrich-Schiller-University Jena, Jena, Germany

[3] Quadram Institute Bioscience, Norwich Research Park, Norwich, UK

[4] National Genomics Data Center, China National Center for Bioinformation, Beijing Institute of Genomics, Chinese Academy of Sciences, Beijing, China

[5] University of Chinese Academy of Sciences, Beijing, China

[6] Department of Molecular Virology, Faculty of Biology, Adam Mickiewicz University, Poznan, Poland

[7] Department of Microbiology, Ohio State University, Columbus, OH, USA

[8] Center of Microbiome Science, Ohio State University, Columbus, OH, USA

[9] EMERGE Biology Integration Institute, Ohio State University, Columbus, OH, USA

[10] Adelaide Medical School, Faculty of Health and Medical Sciences, The University of Adelaide, Adelaide, South Australia, Australia.

[11] Department of Surgery - Otolaryngology Head and Neck Surgery, University of Adelaide and the Basil Hetzel Institute for Translational Health Research, Central Adelaide Local Health Network, South Australia, Australia.

[12] National Center for Biotechnology Information, National Library of Medicine, National Institutes of Health, Bethesda, MD, USA.

[13] Department of Population Health and Reproduction, University of California, Davis, CA, USA

[14] DOE Joint Genome Institute, Lawrence Berkeley National Laboratory, Berkeley, CA, USA

[15] Department of Biochemistry, Institute of Chemistry, University of São Paulo, São Paulo, SP, Brazil

[16] Department of Microbiology, Immunology and Transplantation, Laboratory of Viral Metagenomics, KU Leuven, Leuven, Belgium

[17] Faculty of Automatic Control, Electronics and Computer Science, Silesian University of Technology, Gliwice, Poland

[18] Unaffiliated (independent scientist)

[19] Flinders Accelerator for Microbiome Exploration, Flinders University, Bedford Park, SA, Australia

[20] Institute of Science and Technology for Brain-Inspired Intelligence and MOE Frontiers Center for Brain Science, Fudan University, Shanghai, China
[21] Instituto de Ciências Biomédicas, Universidade de São Paulo, São Paulon, Brazil
[22] Theoretical Biology and Bioinformatics, Science4Life, Utrecht University, Utrecht, the Netherlands
[23] Key Laboratory of South China Sea Fishery Resources Exploitation & Utilization, Ministry of Agriculture and Rural Affairs, South China Sea Fisheries Research Institute, Chinese Academy of Fishery Sciences, Guangdong, China
[24] Institute of Bioinformatics, University Medicine Greifswald, Germany
[25] School of Biological Sciences, Seoul National University, Seoul, Republic of Korea
[26] Interdisciplinary Program in Bioinformatics, Seoul National University, Seoul, Republic of Korea
[27] Institut Pasteur, Université Paris Cité, Cell Biology and Virology of Archaea Unit, Paris, France
[28] Frederick, Maryland, USA
[29] Department of Microbiology, University of Alabama at Birmingham, Birmingham, AL, USA
[30] Institut de Recherche sur la Biologie de l'Insecte, UMR 7261 CNRS-Université de Tours, Tours, France
[31] Department of Computer Science, City University of Hong Kong, Hong Kong SAR, China
[32] Department of Marine Microbiology and Biogeochemistry, NIOZ Royal Netherlands Institute for Sea Research, Texel, The Netherlands
[33] Texas A&M University, Department of Entomology, College Station, TX, USA
[34] Aero-Aquatic Virus Research Group, Faculty of Mathematics and Computer Science, Friedrich Schiller University Jena, Germany
[35] Centre for Ecology and Evolution in Microbial Model Systems (EEMiS), Department of Biology and Environmental Science, Linnaeus University, Kalmar, Sweden
[36] Leibniz Institute on Aging - Fritz Lipmann Institute (FLI), Jena, Germany
[37] Department of Biological Sciences, Universidad de los Andes, Bogotá, Colombia
[38] Consiglio Nazionale delle Ricerche, Istituto per la Protezione Sostenibile delle Piante, Sede Secondaria di Bari, Bari, Italy
[39] Department of Agricultural Science and Plant Protection, Mississippi State University, Mississippi State, MS, USA
[40] Institute for Genomics, Biocomputing and Biotechnology, Mississippi State University, Mississippi State, MS, USA
[41] Department of Information Engineering, Chinese University of Hong Kong, Hong Kong SAR, China
[42] Institute of Biomedicine, University of Turku, Turku, Finland
[43] Nuffield Department of Medicine, University of Oxford, Oxford, UK
[44] Institute of Molecular Biology and Genetics, Seoul National University, Seoul, Republic of Korea
[45] Artificial Intelligence Institute, Seoul National University, Seoul, Republic of Korea
[46] Department of Electrical Engineering, City University of Hong Kong, Hong Kong SAR, China
[47] BAICSM, State Key Laboratory of Green Biomanufacturing, College of Life Science and Technology, Beijing University of Chemical Technology, Beijing, China

[48] Melbourne Data Analytics Platform, The University of Melbourne, Parkville, VIC, Australia
[49] School of Applied Sciences, College of Health, Science and Society, University of the West of England, Bristol, UK
[50] The Biodesign Center for Fundamental and Applied Microbiomics, School of Life Sciences, Arizona State University, Center of Evolution and Medicine, Tempe, AZ, USA
[51] Structural Biology Research Unit, Department of Integrative Biomedical Sciences, University of Cape Town, Cape Town, South Africa
[52] School of Mathematical Sciences and LPMC, Nankai University, Tianjin, China.
[53] Department of Microbiology and Immunology at The Peter Doherty Institute for Infection and Immunity, The University of Melbourne, Melbourne, Victoria, Australia
[54] Centre for Pathogen Genomics, University of Melbourne, Victoria, Australia
[55] School of Life Sciences and Biopharmaceutics, Guangdong Pharmaceutical University, Guangdong, China
[56] Department of Anaesthesia and Intensive Care, Faculty of Medicine, The Chinese University of Hong Kong, Hong Kong SAR, China
[57] Li Ka Shing Institute of Health Sciences, The Chinese University of Hong Kong, Hong Kong SAR, China
[58] Computational Biology Research Unit, Faculty of Biology, Adam Mickiewicz University, Poznan, Poland
