## Supplementary figures and images for "First community challenge for automated virus taxonomy"

### Supplementary figure 1

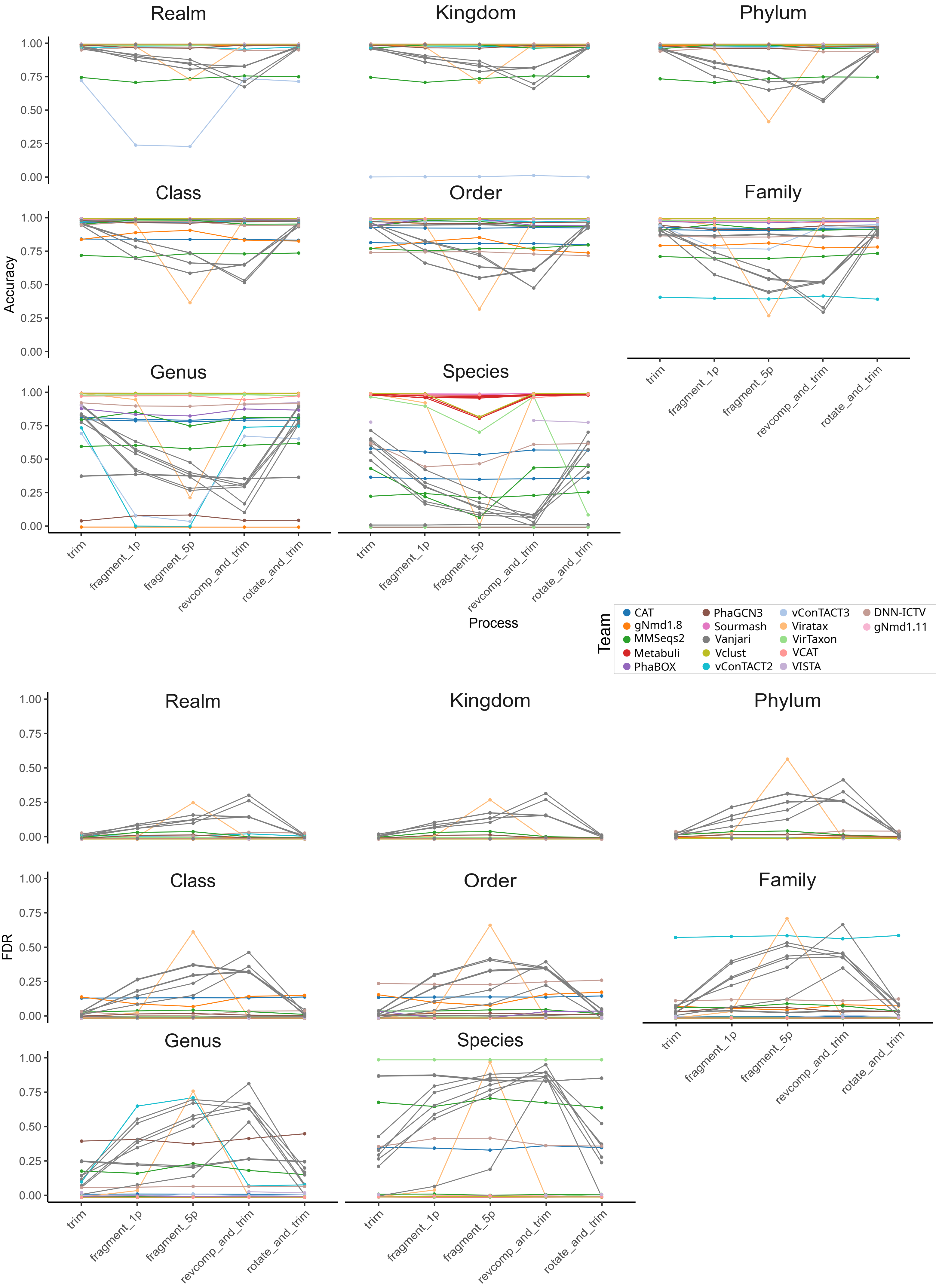

### Supplementary figure 2

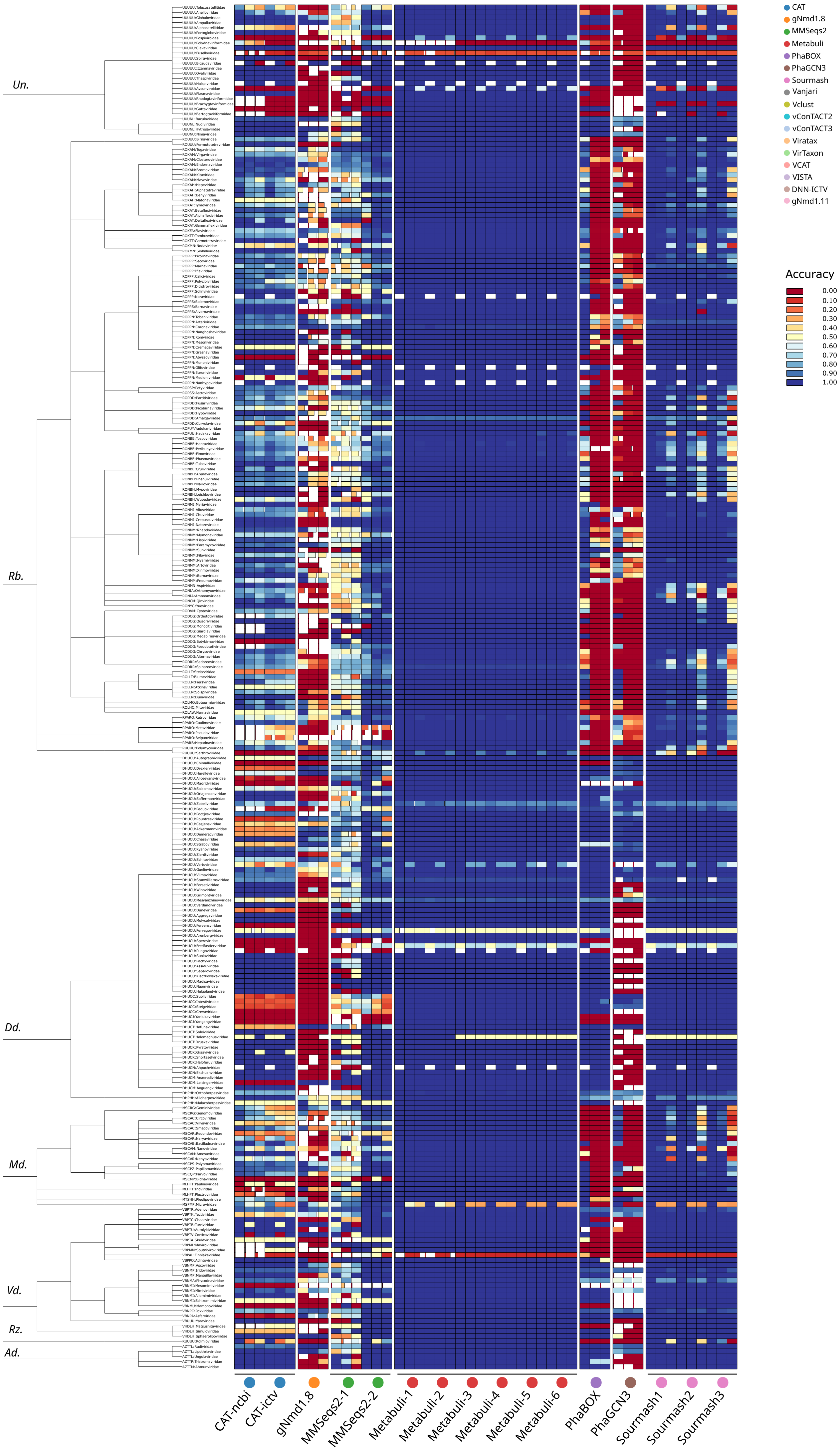

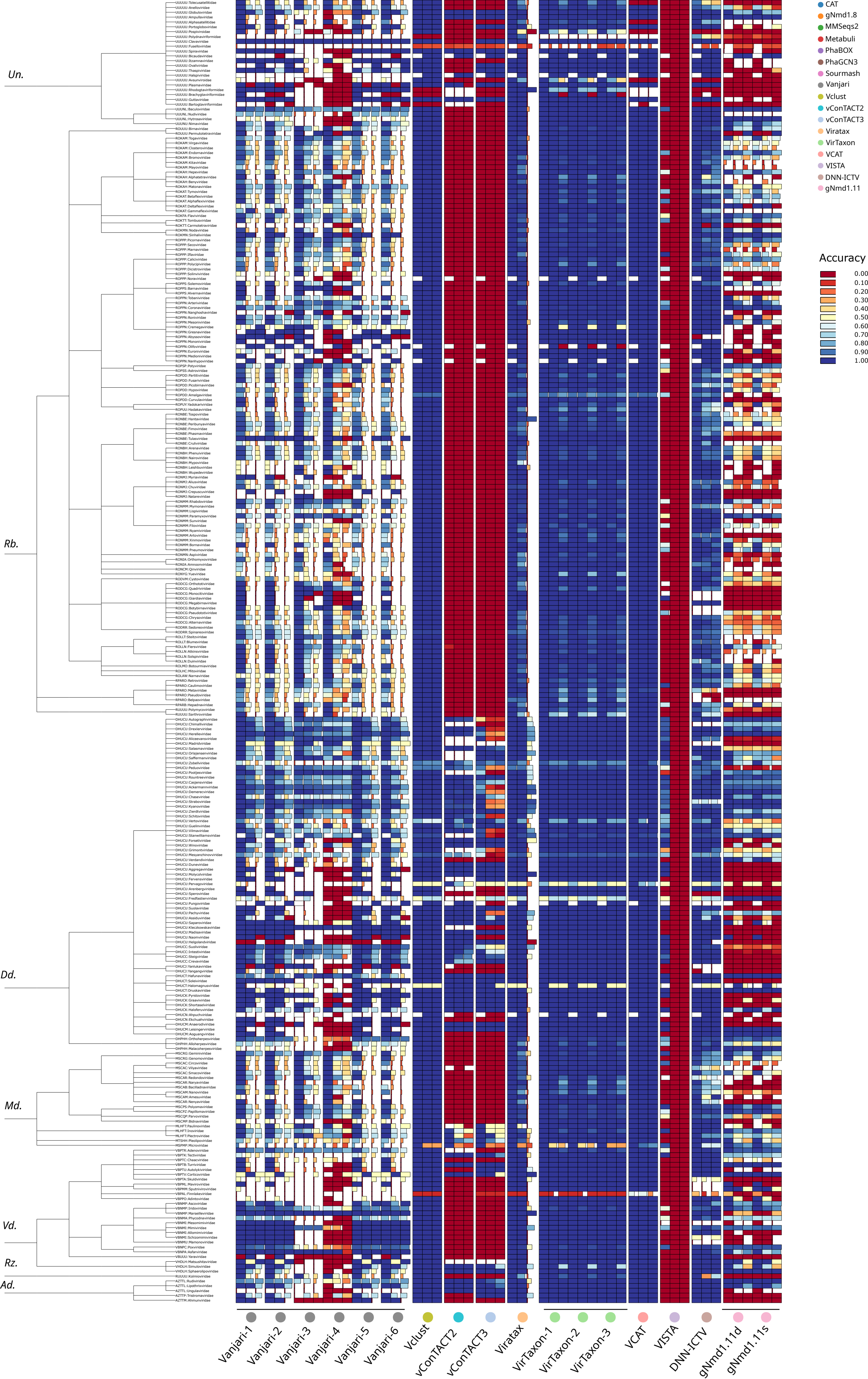

### Supplementary figure 3

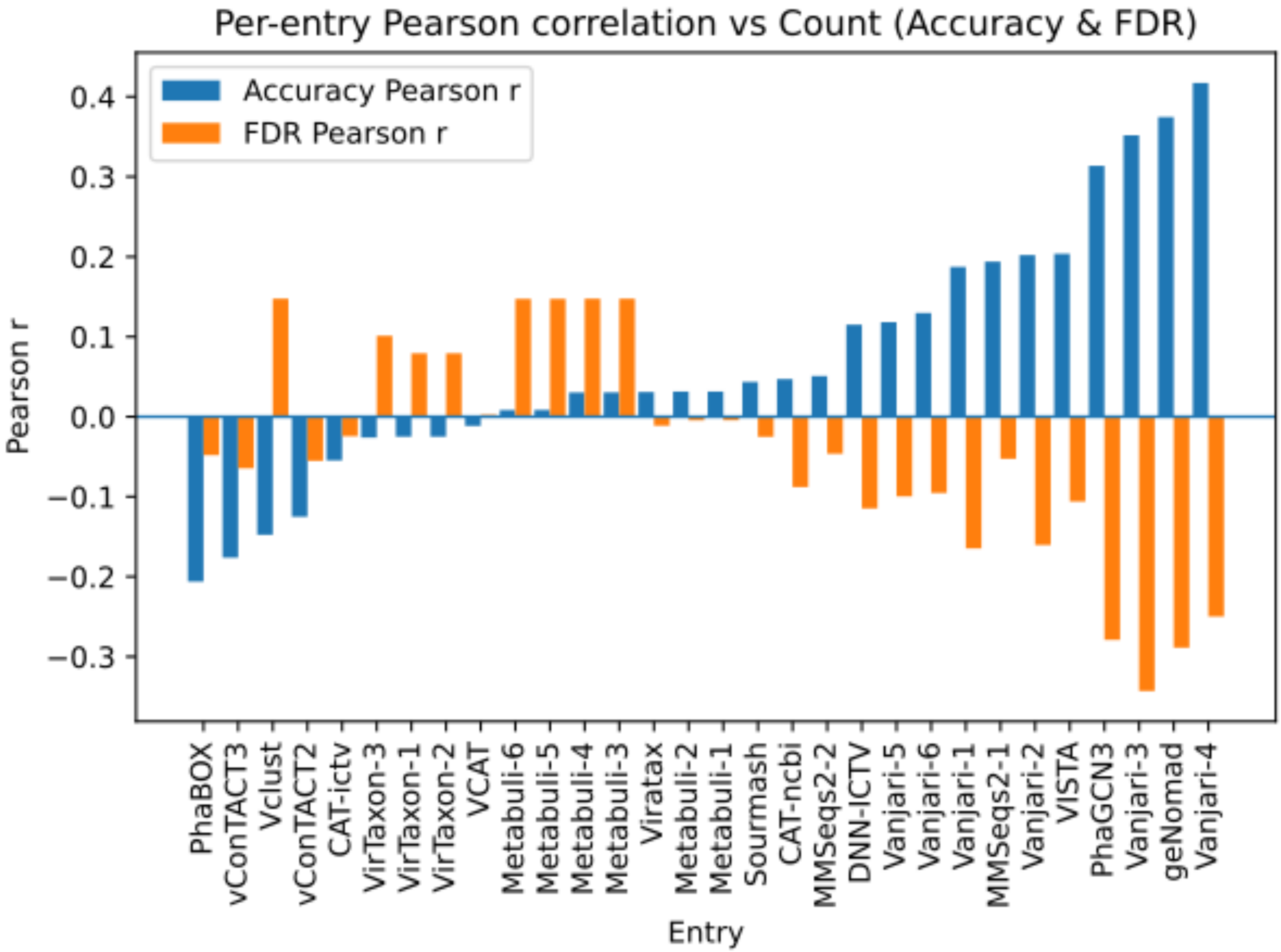

### Supplementary figure 4

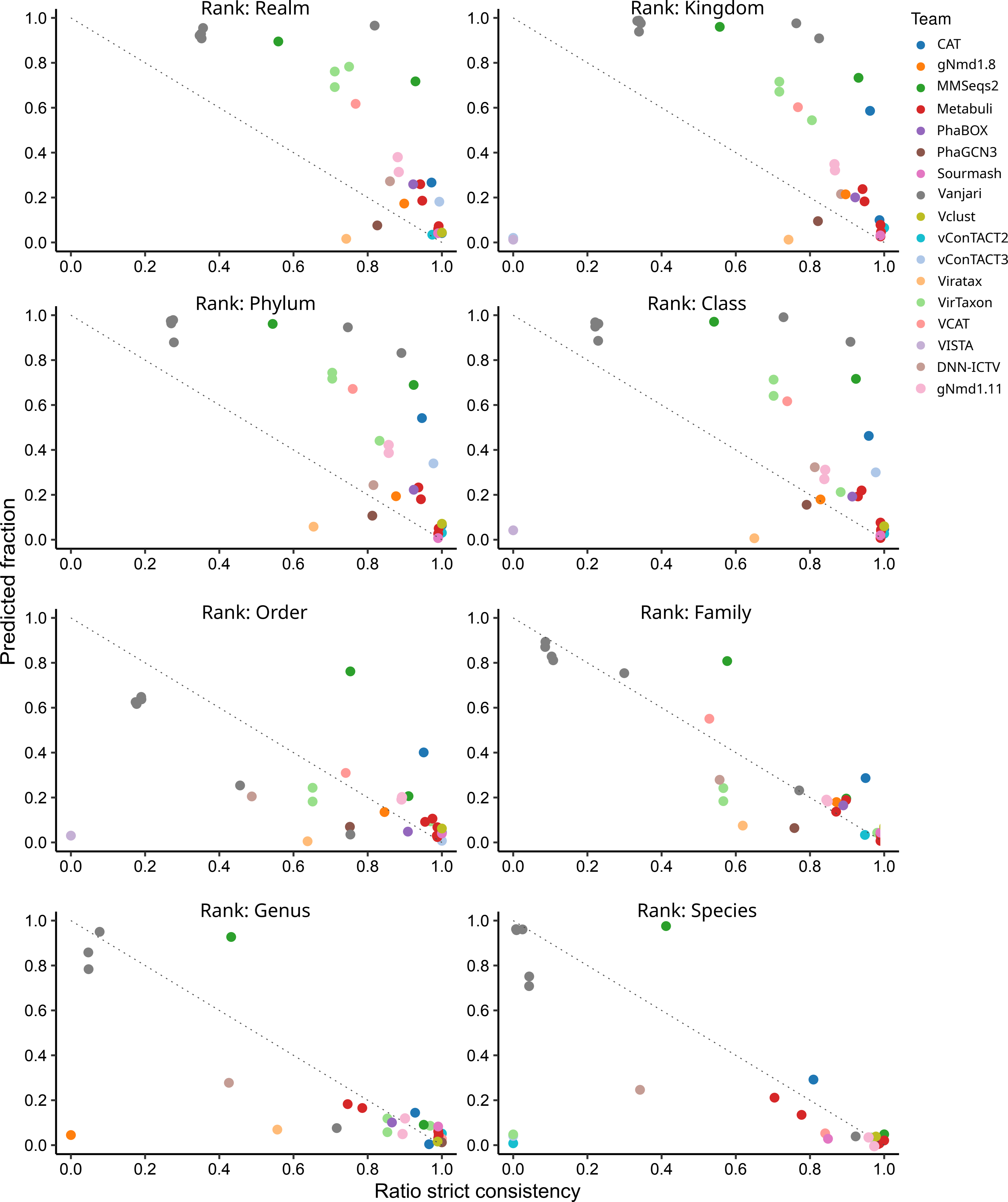

### Supplementary figure 5

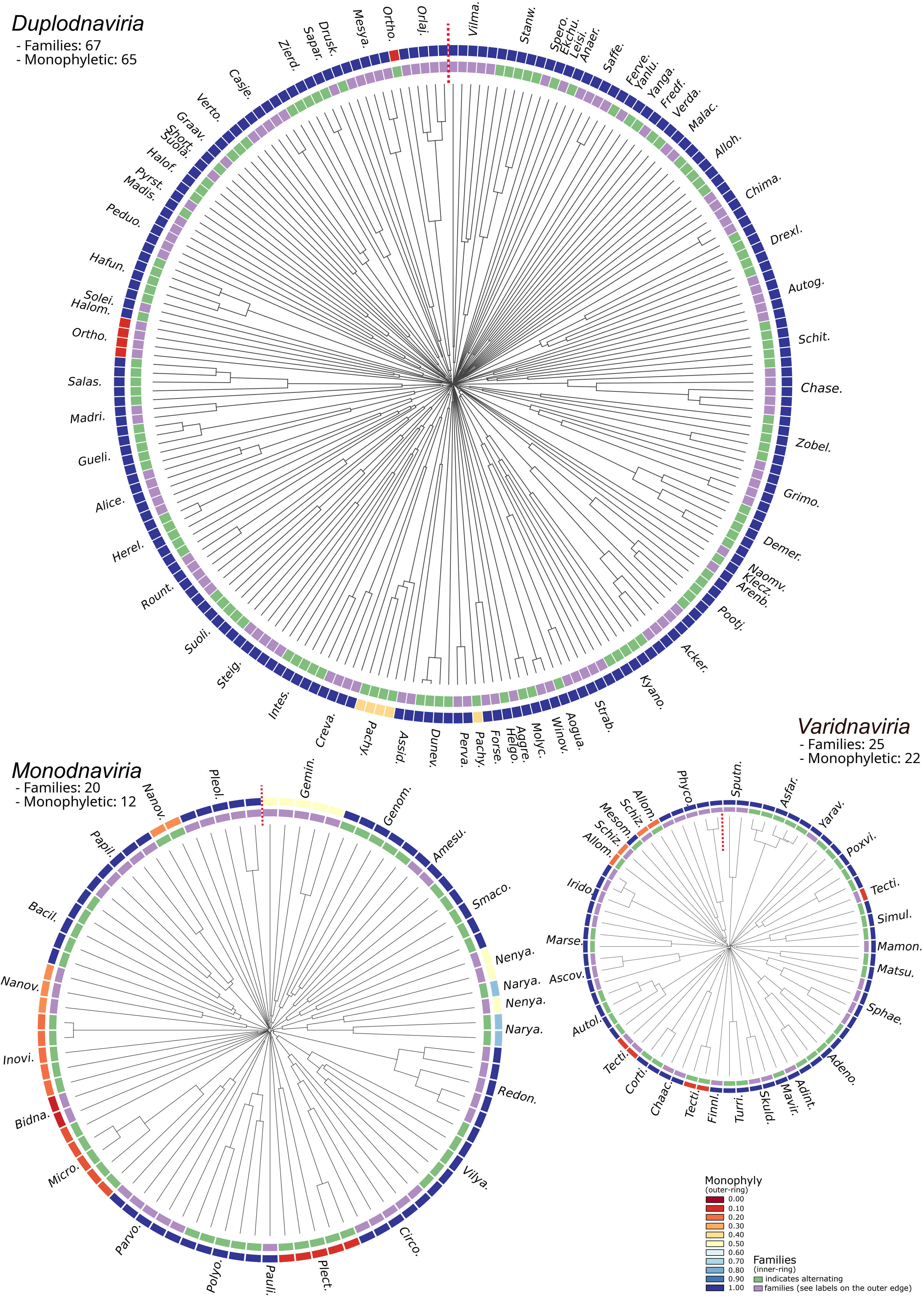

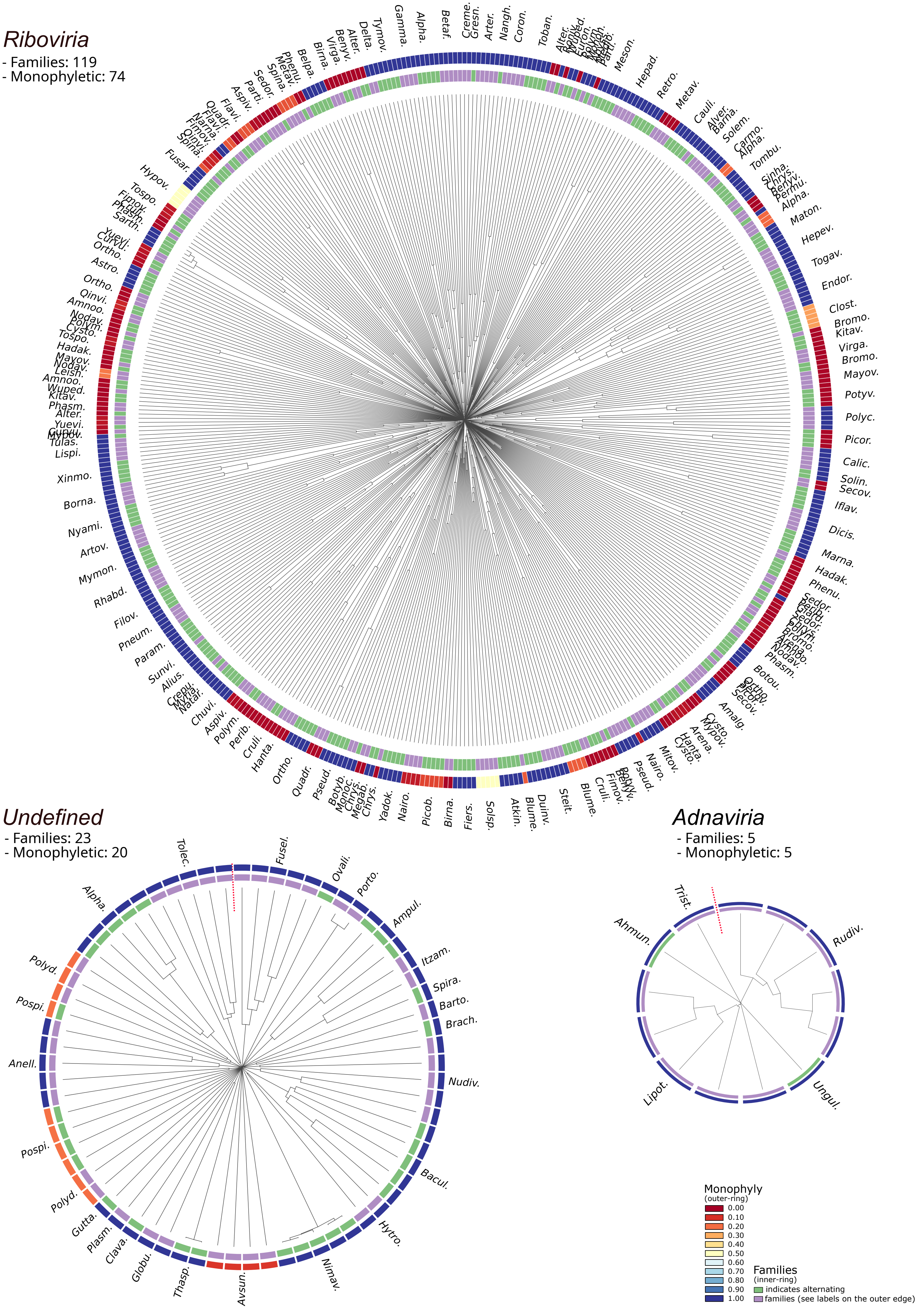

### Supplementary figure 6

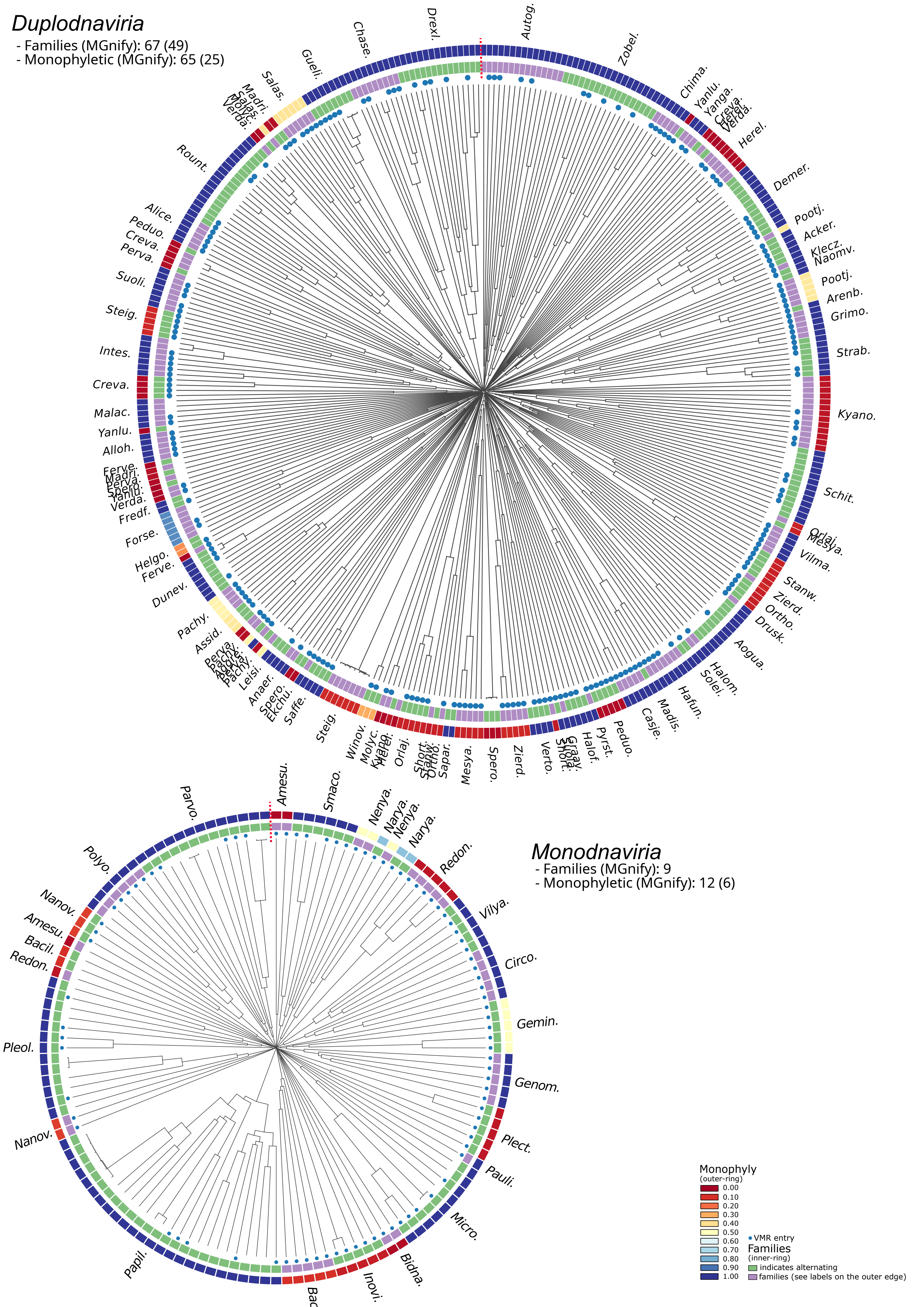

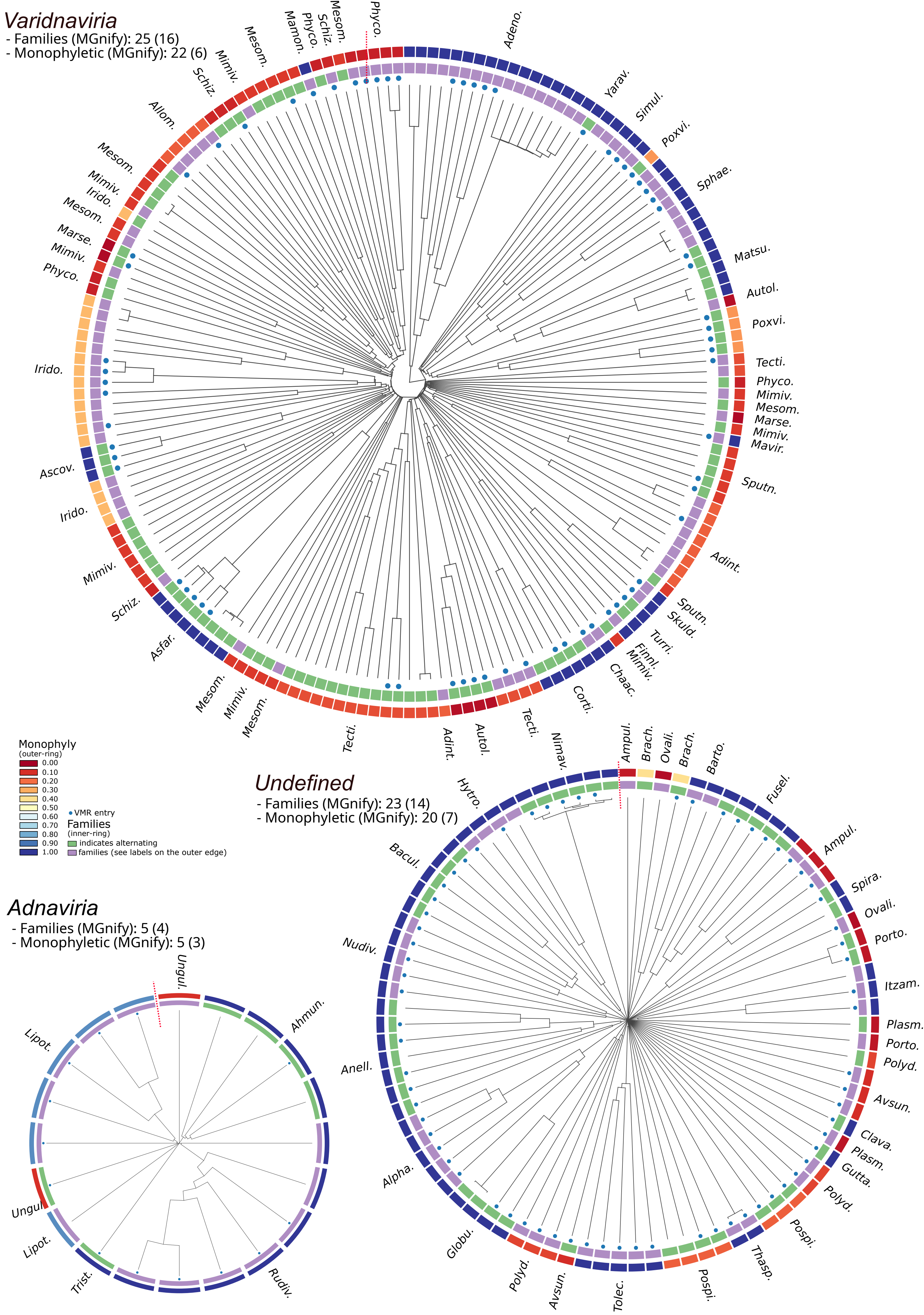

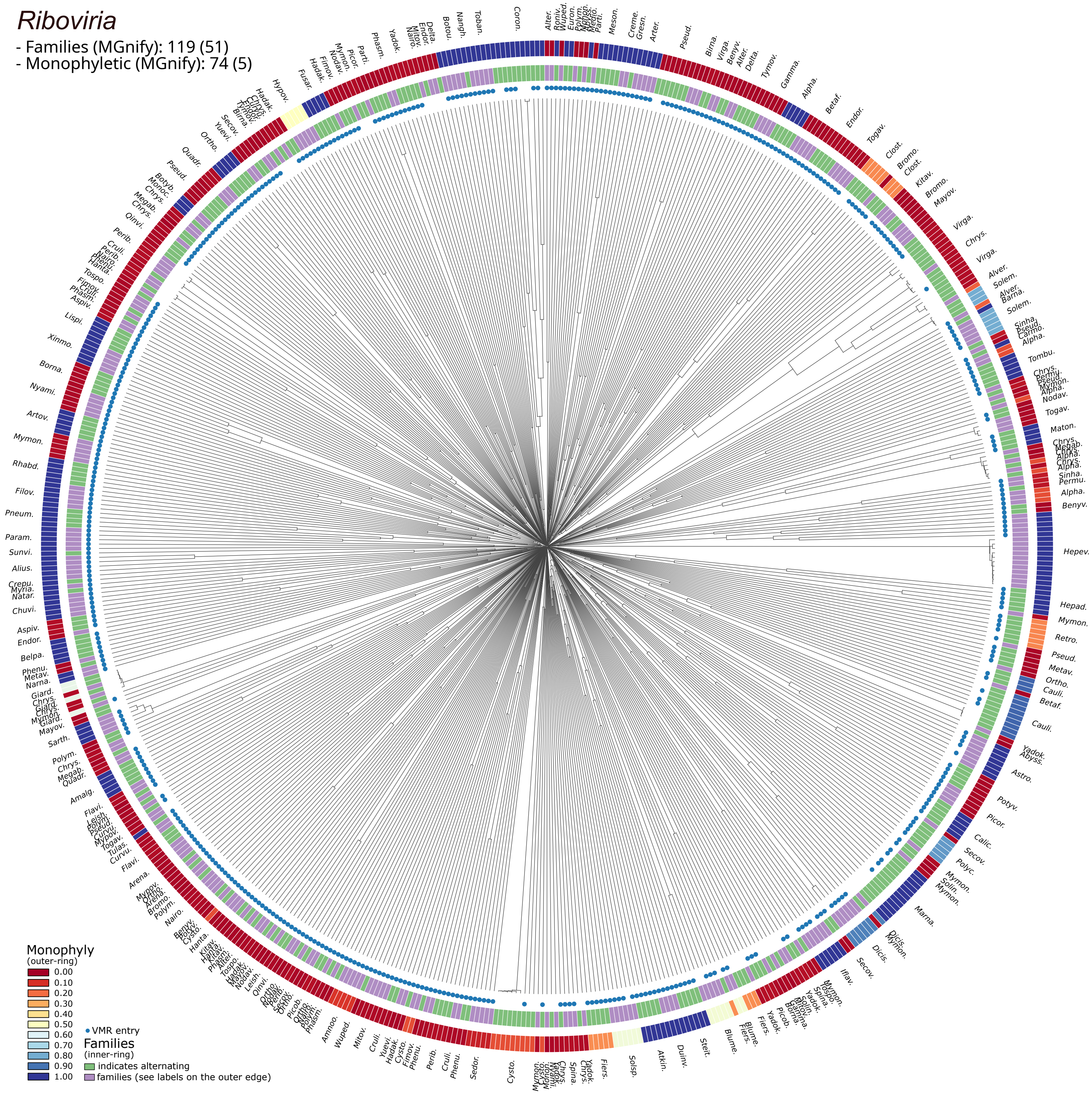
